# High-frequency oscillations and replay in a two-population model of hippocampal region CA1

**DOI:** 10.1101/2021.06.08.447523

**Authors:** Wilhelm Braun, Raoul-Martin Memmesheimer

## Abstract

Hippocampal sharp wave/ripple oscillations are a prominent pattern of collective activity, which consists of a strong overall increase of activity with onmodulated (140 – 200 Hz) ripple oscillations. Despite its prominence and its experimentally demonstrated importance for memory consolidation, the mechanisms underlying its generation are to date not understood. Several models assume that recurrent networks of inhibitory cells alone can explain the generation and main characteristics of the ripple oscillations. Recent experiments, however, indicate that in addition to inhibitory basket cells, the pattern requires *in vivo* the activity of the local population of excitatory pyramidal cells. Here we study a model for networks in the hippocampal region CA1 incorporating such a local excitatory population of pyramidal neurons and investigate its ability to generate ripple oscillations using extensive simulations. We find that with biologically plausible values for single neuron, synapse and connectivity parameters, random connectivity and absent strong feedforward drive to the inhibitory population, oscillation patterns similar to *in vivo* sharp wave/ripples can only be generated if excitatory cell spiking is triggered by short pulses of external excitation. Specifically, whereas temporally broad excitation can lead to high-frequency oscillations in the ripple range, sparse pyramidal cell activity is only obtained with pulse-like external CA3 excitation. Further simulations indicate that such short pulses could originate from dendritic spikes in the apical or basal dendrites of CA1 pyramidal cells, which are triggered by coincident spike arrivals from hippocampal region CA3. Finally we show that replay of sequences by pyramidal neurons and ripple oscillations can arise intrinsically in CA1 due to structured connectivity that gives rise to alternating excitatory pulse and inhibitory gap coding; the latter implies phases of silence in specific basket cell groups and selective disinhibition of groups of pyramidal neurons. This general mechanism for sequence generation leads to sparse pyramidal cell and dense basket cell spiking, does not rely on synfire chain-like feedforward excitation and may be relevant for other brain regions as well.

**Author summary:** During certain phases of sleep, rest and consummatory behavior the hippocampus brain area of many species, including humans, is known to intermittently generate strong high frequency oscillations. These oscillations are important for memory formation and consolidation. To date, the mechanisms underlying their generation remain incompletely understood. We find that in unstructured networks carefully designing how excitation is transmitted in the hippocampus is required for the generation of robust fast oscillations in its main output region. Broad, temporally extended excitation of cells results in unrealistic single cell activity, whereas temporally narrow input that differs from cell to cell gives rise to oscillations with realistic single cell and network activity. We show that the biophysical mechanism to generate the required temporally narrow excitation may be related to spiking events in the dendrites, which are triggered by coincident input. Our results in structured networks suggest that the interplay of hippocampal excitation and inhibition can serve as a means to generate robust sequential activity, which is thought to be crucial for memory formation and recall. The sequence generation mechanism also leads to strong high frequency oscillations with sparse excitatory cell and frequent inhibitory cell spiking, as observed in the hippocampus.

## Introduction

Hippocampal sharp wave/ripples (SPW/Rs) are remarkable both from a neurophysiological viewpoint and in their behavioral impact: On the one hand, they consist of strong increases of spiking activity in large parts of local neuron populations (the sharp wave) together with oscillatory, extraordinarily coherent neuronal discharges (ripples); on the other hand, SPW/Rs have directly been shown to be important for memory consolidation [1–6] and might be involved in planning of future actions [1, 7]. Perhaps the most striking signature of memory consolidation is the phenomenon of hippocampal replay (see [8] for a recent review) during which sequences of pyramidal cell action potentials encoding location, so-called place cells [9], are repeated on a faster timescale during SPW/R complexes.

SPW/Rs occur in all mammalian species that have been investigated for them, including humans [1, 10]. Similar activity has been observed in fish [1], reptiles [11] and birds [12], indicating that the brain circuits involved in their generation are evolutionary old. The existence of SPW/Rs in non-mammalian species remains, however, a controversial subject [1, 12].

Our understanding of the role of SPW/Rs in the brain would benefit from a detailed understanding of the underlying generative mechanism, which includes the external excitation, the number and identity of the involved neurons, their interaction and the required pattern of connectivity.

SPW/Rs are most prominently observed in the hippocampal region CA1. The main source of external excitation delivered to CA1 during a SPW/R event is a sharp, wave-like depolarization from area CA3 delivered via the Schaffer collaterals (SCs) to the basal and proximal apical dendrites of CA1 pyramidal cells, which are located in the CA1 strata oriens and radiatum, respectively [1, 13–15]. This input likely generates the sharp wave of spiking activity in CA1. We note that in *in vitro* experiments, SPW/Rs can also be generated in the isolated region CA1 [16–18]. A typical sharp wave has a duration of 40 – 100 ms [1]. The co-occurring ripple oscillations were first observed as oscillations in the local field potential (LFP) when recording in the CA1 stratum pyramidale [13, 14, 19]. Direct measurements of spiking activity and modeling studies show that these LFP ripples are mainly caused by the temporally precise, sparse firing of excitatory pyramidal cells [20–22], possibly with a smaller contribution from interneuronal activity [1, 22]; ref. [23] found in CA3 slices that GABAergic interneurons are both necessary and sufficient to generate ripple oscillations in the LFP, even when excitation is blocked.

The onmodulated CA1 ripples are uncorrelated with weaker, lower frequency ripples occurring in CA3 and are thus considered to be generated by CA1 networks [14, 24]. How CA1 generates the fast ripple oscillation is to date not clear; the focus of this article is to design minimal CA1 network models that incorporate plausible CA1 connectivity and produce SPW/Rs with realistic network and single-cell activity.

Several classes of competing models exist for ripple generation. The first class, which we term *interneuron ripples* (IR) (narrowing down the definition of inhibition-first ripples given in [25]), are models where mainly the inhibitory neurons generate the rhythm and entrain the excitatory ones. The standard, network model of this class (*interneuron network ripples*, INR) posits that the ripples in CA1 are generated by tonic excitation of a recurrent network of fast-spiking basket cells (BCs), usually of the parvalbumin-positive (PV+) immunoreactive type [23, 25–31]. The oscillation unfolds as follows, according to a mechanism also thought to underlie certain gamma oscillations (interneuron network gamma, ING) [32–36]: A peak of interneuron spiking activity induces a peak in recurrent inhibition after a short delay, which leads to a subsequent valley in inhibitory spiking activity. The ensuing lack of recurrent inhibition allows for a subsequent activity peak and so on. In support of this mechanism, theoretical studies have shown that fast oscillations in the ripple range can be generated easily in presence of short synaptic time constants for the recurrent GABAergic inhibitory currents [27, 37], both in integrate-and-fire (IF) [37] and more detailed Hodgkin-Huxley conductance-based neuron models [27]. Such models have recently been successfully applied to explain phenomena such as intraripple frequency accommodation and the dependency of the network frequency on GABA modulators [25]. Another model that falls within the IR class proposes that a single neuron mechanism underlies the ripple rhythm: calcium spikes in the dendrites of PV+BCs generate high-frequency oscillations in the membrane potential and output action potentials occur preferentially at the oscillation peaks [38].

Recently it has become clear, however, that an IR model might be insufficient for the description of the ripple mechanism. *In vivo* experiments [22, 39] might suggest a crucial involvement of the local pyramidal cell population in ripple generation. It was shown that optogenetic excitation of a small number of excitatory (E) pyramidal cells in CA1 can generate high-frequency (ripple-like) oscillations (HFOs) and prolongs SPW/R events. In contrast, optogenetic activation of inhibitory (I) interneurons did not generate ripples and stopped them during SPW/Rs events; some coherent spiking activity in the ripple range could, however, be generated. Further, I cell spiking, caused mainly by excitatory-to-inhibitory connections, was necessary to stabilize the ripple oscillations. It was thus proposed that interactions between E and I cell populations after external excitation of E cells are both necessary and sufficient to generate ripple oscillations *in vivo* [22]. These results suggest a second class of models, which are the focus of the present article and which may be called *pyramidal interneuron ripples* (PIRs). In them, both populations are similarly important for rhythm generation. In the standard, network model of this class (*pyramidal interneuron network ripples* (PINRs)), the ripple oscillation is generated by an interplay of the two recurrently connected cell populations, where the excitatory population receives external drive. The excitatory-inhibitory loop then generates an oscillation according to a mechanism that is called pyramidal interneuron network gamma (PING) in the context of gamma oscillations [33–36]: The excitatory population excites the inhibitory population, which in turn transiently reduces E cell activity. After enough excitation has build up in the E cell network, the cycle restarts. If the interneuron network alone can already oscillate due to its recurrent inhibition, adding an excitatory population yields, for weak synchrony, a compromise between the oscillations that emerge due to the two mechanisms [27, 37] or, for strong synchrony, competition between them [36, 40]. Networks of the former, weakly synchronized type have been suggested as model for ripple oscillations. Adding the excitatory population usually decreases the oscillation frequency of the network, but can also increase it [27, 37].

In the last class of models, which we term *pyramidal ripple* (PR) models (equivalent to the excitation-first class in [25]), the excitatory neuron population mainly governs the ripple rhythm [41–44] and entrains the inhibitory neurons by local excitatory-to-inhibitory synaptic connections. The presence of inhibition can nevertheless be crucial, as it may sharpen the ripples and prevent unrhythmic pathological spiking activity [43].

The first subclass of PR models posits that gap junctions between the axons of pyramidal cells enable spike propagation in the axonal plexus. Through antidromic invasion into the somata, these generate pulses of somatic spikes at ripple frequency [16, 18, 41, 42, 45, 46]. While there is no concluding, ultrastructural evidence for axo-axonic gap junctions between CA1 pyramidal cells [47], ref. [45] showed that axons of CA1 pyramidal neurons are dye coupled. Further, gap junctions have been ultrastructurally demonstrated between dendritic and somatic locations in CA1 pyramidal neurons [48] and between axons of dentate gyrus principle cells [49]. Experiments, however, indicate that somatic spikes are generated by orthodromic excitation during SPW/Rs *in vivo* [50].

The second subclass of PR models proposes that synchrony propagation supported by nonlinear dendrites underlies SPW/Rs [43, 44, 51–55]. Specifically, it assumes that spikes in the basal dendrites of CA1 pyramidal neurons are generated by sufficiently synchronous recurrent excitation [43, 44]. Whether the recurrent excitatory connectivity in CA1, which is very sparse (typical values for coupling probabilities are between 1 and 2% [56, 57]), suffices to generate these dendritic spikes, is currently unclear. Models incorporating such non-linearities [43, 44] have successfully reproduced many experimental findings, including sparse firing of pyramidal cells and the phase relation of the excitatory and inhibitory cell population.

A successful model for SPW/R generation must reproduce the main properties of ripple oscillations in CA1. The most important criterion is that the activity of E and I populations should be modulated at a frequency in the ripple range, between 140 and 200 Hz. *In vivo* CA1 ripples lie in the lower (slow wave sleep ripples) and middle (awake ripples) part of this range[1, 20, 22, 58], *in vitro* ripples in the upper part. A model for *in vivo* and *in vitro* ripples must therefore be able to cover the entire frequency range [1, 16, 18].

The second main property of ripple oscillations is that pyramidal neurons fire one to two orders of magnitude less frequently than interneurons [20, 59]. During SPW/Rs, pyramidal cells have firing rates between 0 and 10 Hz, with a bias towards smaller frequencies, so that the mean firing rate during ripples is between 1 and 2 Hz [60]. More concretely, pyramidal cells tend to fire only once during a sharp wave/ripple event [22, 39], which is composed of multiple ripple waves (or: ripple cycles). This entails that different sets of pyramidal neurons are active on each ripple wave. Fast-spiking PV+BCs [61] have much higher average firing rates and often contribute a spike to every ripple wave [13, 20, 39]. Their firing rates during a SPW/Rs typically lie between 10 and 200 Hz [20, 59, 62]. 10 – 20% of pyramidal cells typically discharge during an SPW/R event, whereas 80% and more of interneurons fire[20, 22, 63, 64].

In this paper, we first show that simply augmenting previous inhibition-first models by adding pyramidal neurons is not sufficient to generate realistic single-cell and network dynamics, unless the interneurons receive very strong feedforward excitatory drive. We show that nevertheless the generation of high-frequency oscillations in the right frequency range with realistic single-neuron dynamics is possible with two-population models if excitation to CA1 pyramidal cells is delivered in a temporally inhomogeneous fashion. Such inhomogeneity could be brought about by dendritic action potentials depolarizing the soma of CA1 pyramidal cells, which are mediated by coincidence detection from incoming CA3 spikes. Finally, we show that in such a two-population model, robust sequences of pyramidal cell firing activity, as observed during hippocampal replay, can be generated intrinsically in CA1 by a mechanism involving disinhibition of selected groups of pyramidal cells.

## Materials and methods

We consider three variants, model 1, 2 and 3, of a two-population model consisting of *N_E_* excitatory pyramidal cells (PCs), the principal cells of hippocampal region CA1, and *N_I_* inhibitory parvalbumin-positive BCs (Fig 1).

**Fig 1.**
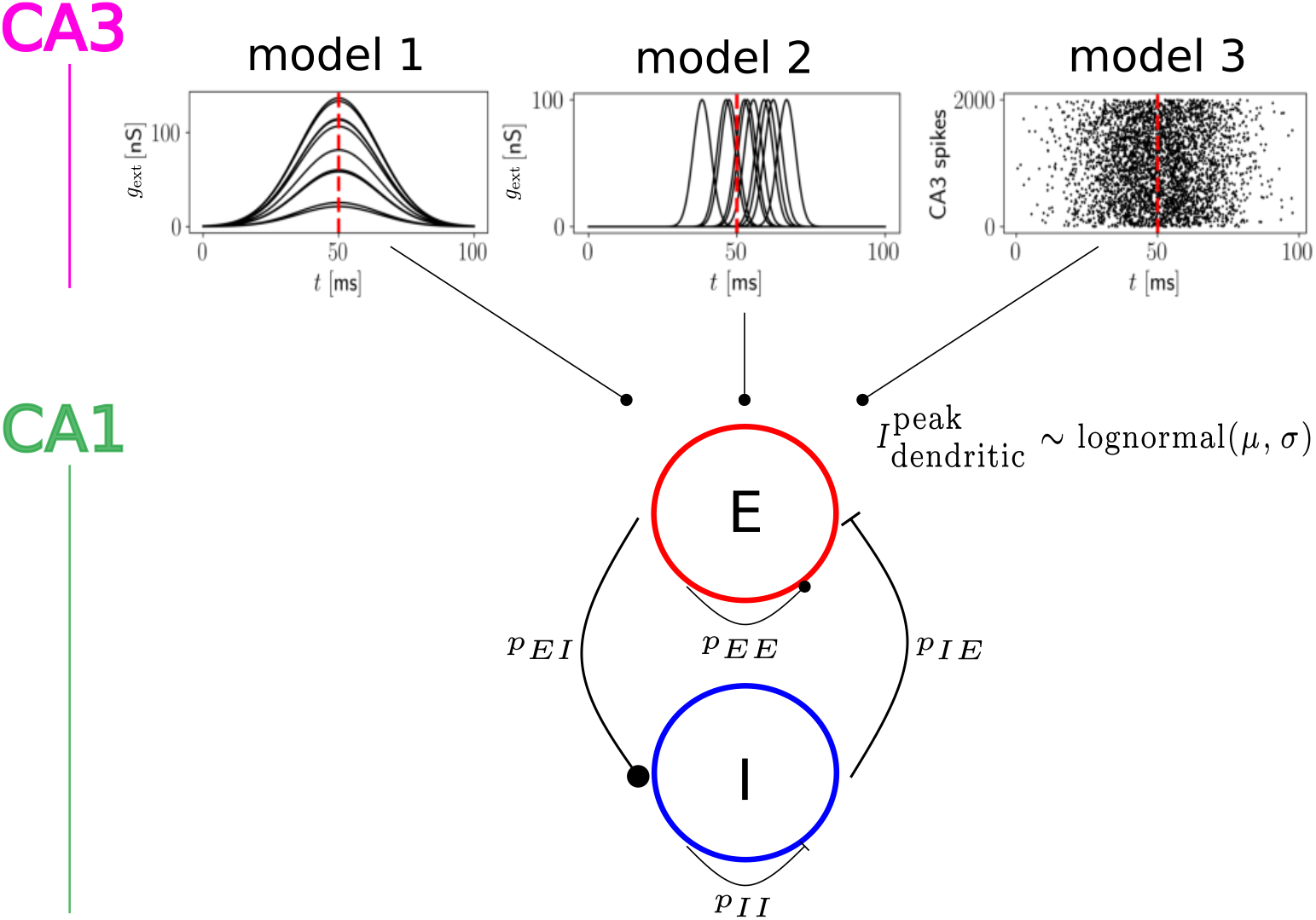
Model overview. Schematic drawing of the three models considered in this paper. They consist of a population of *N_E_* excitatory and a population of *N_I_* inhibitory CA1 neurons, which are connected with probabilities *p_αβ_, α, β ∈ {E, I}* (lower panel) and receive external input from CA3 (upper panel). The models differ in this external input: In model 1 and 2, it is represented by the excitatory conductances *g*_ext_ (Eq 6) that it evokes in *n_E_* CA1 neurons. In model 1, the time courses of these conductances are broad and for each neuron centered around the same time (upper left panel). In model 2, the time courses are short, pulse-like and centered around different times (upper middle panel). In model 3, CA3 is represented by a population of input neurons. Each such neuron fires according to an inhomogeneous Poisson process with Gaussian rate profile given by Eq 7 (upper right panel). Spike transmission from CA3 to CA1 is then filtered by a strong apical or basal dendrite of the receiving CA1 neuron: Besides the linear input transmission, sufficiently coincident input evokes dendritic spikes. These generate somatic currents whose peak strengths follow a lognormal distribution (Eq 2) across neurons.

Their dynamics are governed by a conductance-based integrate-and-fire neuron model. It reads for a neuron in the excitatory population

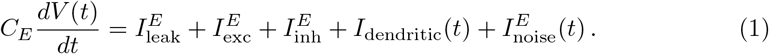

The leak current is given by 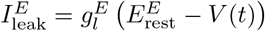 and the excitatory and inhibitory currents are given by 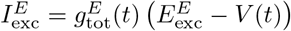 and 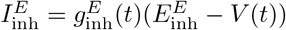, respectively. The total excitatory conductance consists of two parts: 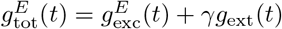 representing input from recurrent excitatory synapses and external input. The external input is a direct, not dendritically amplified, input present in model 1 and 2. It models input from CA3 or an optogenetic stimulation delivered in experiments [22]. *γ* ∈ {0, 1} determines whether this drive is present in the model, i.e. *γ* = 1 in m els 1 and 2, whereas *γ* = 0 in model 3. The noise current 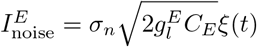 is a Gaussian white noise input, independent between neurons and with strength *σ_n_* = 1 mV.

*I*_dendritic_ is a somatic current triggered by apical or basal dendritic spikes and only present in model 3. We implement the generation of dendritic spikes via a temporal coincidence detection mechanism [43, 44]: As soon as the dendrite receives sufficiently many spikes from its presynaptic CA3 neurons within the dendritic integration window *w_D_* = 2 ms (< 3 ms [65, 66]), a fast and strong, but not necessarily suprathreshold excitatory current *I*_dendritic_ is generated. It depolarizes the somatic membrane potential a delay *τ_D_* = 2 ms after the coincidence was detected. There is a refractory period *t_r,D_* = 5.0 ms during which no further dendritic spikes are triggered – this means that after a dendritic spike, there cannot be another one for a duration of *t_r,D_*. We checked that our results do not depend critically on the exact value of *t_r,D_* by performing simulations with smaller (*t_r,D_* = 2 ms) and larger (*t_r,D_* = 10 ms) values. The results that we will report below for model 3 remained qualitatively unchanged, only the parameters of the lognormal distribution (Eq 2) where HFOs in the ripple range occur shifted such that a smaller (larger) *t_r,D_* was compensated for by an increase (decrease) in *μ* and/or *σ*.

We assume that every pyramidal neuron has one dendritic compartment where it receives the relevant input from CA3. This models the single main apical dendrite in CA1 pyramidal neurons [15, 65, 67] and its several oblique ones emanating from it. We do not explicitly model the compartment in this study, but instead focus on the impact a dendritic spike has on the somatic membrane voltage. All inputs from CA3 contribute in the same way to dendritic spike generation. In particular we do not separately model local effects such as the generation of weak dendritic spikes in the oblique dendrites [68]. Because innervation by CA3 Schaffer collaterals occurs on both basal and apical dendrites of CA1 pyramidal neurons [15], the modeled dendrite may alternatively be interpreted as an influential, privileged basal one [69] (neglecting the contribution of recurrent CA1 excitatory inputs to dendritic spike generation in such a dendrite – dendritic spikes in our model 3 are generated by feedforward Schaffer collateral input). Experiments show that the impact of dendritic spikes on the soma can be different depending on the generating dendrite, but also on the depolarization and the somatic firing history [65, 70, 71]. If not mentioned otherwise we thus assume that the strengths of dendritic spikes across neurons have a lognormal distribution as experimentally found for several other neuronal properties [60, 72]. Specifically the peaks of the currents induced in the soma have a lognormal distribution across neurons,

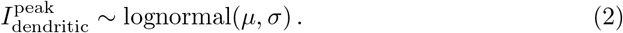

The current in the soma induced by the dendritic spike has a rise time of 1 ms and a decay time of 4 ms [65]. If *V* starts at its resting potential and noise and further input are absent (*σ_n_* = 0 in Eq 1), the minimal peak current to generate a somatic action potential is approximately 1.34 nA. Stronger dendritic spikes can generate multiple somatic spikes: without further inputs and if its peak current is at 10 nA, a dendritic spike generates four somatic ones. However, more than 99.8% of the dendrites in our model 3 simulations without replay have a smaller impact for representative values of *μ* = 0 nA and *σ* = 0.75 nA (see Fig 8 in Results). This follows from the cumulative density function 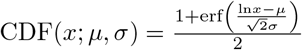 of the lognormal distribution, which yields CDF(*x*; *μ, σ*) = 0.9983 with *x* = 10 nA, *μ* = 0 nA and *σ* = 0.75 nA. We note that somatic action potential bursts in CA1 pyramidal cells are usually generated by a more involved mechanism, where calcium spikes in the apical tuft are generated with the support of backpropagating action potentials [73, 74]. Fig 2 shows examples of dendritic currents and the associated somatic membrane voltage time courses in our model.

**Fig 2.**
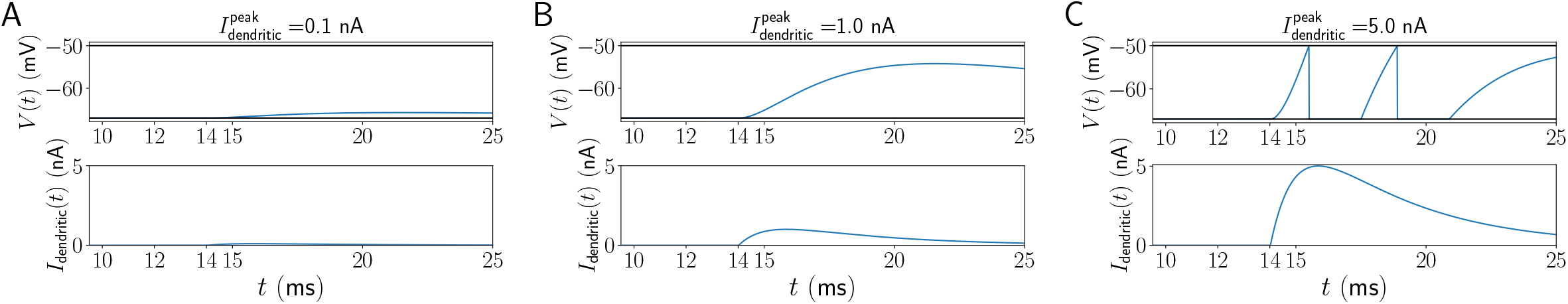
The impact of dendritic spikes on the soma. A dendritic spike is generated at *t* = 12 ms. It is modeled by the dendritic current *I*_dendritic_ (lower subpanels in A,B,C) arriving at the soma *τ_D_* = 2 ms later (here at 14 ms). For small peak dendritic currents 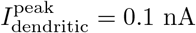, panel A), the somatic depolarization (upper subpanel) is small (here: 1.28 mV) when starting at rest. For larger peak dendritic currents, the somatic depolarization increases (12.78 mV for 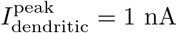, panel B), but remains subthreshold. For 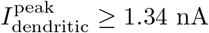, at least one somatic spike is generated; for 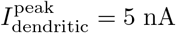 (panel C), two spikes are generated and there is significant depolarization visible in the membrane voltage even after the second spike. The scale for the bottom panels is fixed to facilitate comparison between the different values for the dendritic peak current.

The dynamics of the inhibitory neurons are given by

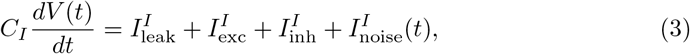

similar to Eq 1. The leak, excitatory and inhibitory currents are 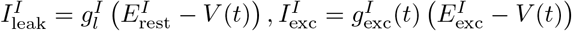 and 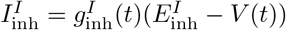 respectively. 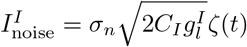 is a Gaussian white noise input, independent between neurons and with strength *σ_n_* = 1 mV. In both the E and I populations, once a neuron reaches its firing threshold 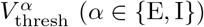 it is reset to 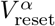 and remains at this potential during an absolute refractory period 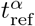.

The neurons are connected by chemical synapses, see next section for details. Gap junctions between inhibitory PV+BCs in CA1 are frequent [47, 75], but their importance for ripple oscillations *in vivo* is unclear [76, 77]. In the context of IR models, recent theoretical results show that adding gap junctions between inhibitory cells has beneficial effects for ripple oscillations, as they enhance synchrony and decrease the minimal number of cells required for a ripple oscillation [78]. Since our models do not require gap junctions and because of their unclear importance for SPW/Rs, we do not consider them in this paper.

We have chosen the single neuron and network parameters in agreement with neuroanatomical and neurophysiological experimental knowledge on the hippocampal area CA1. The data we refer to come mostly from rats, but also from mice [57]. The parameters can be found in the Supporting Information (SI), section Parameters.

### Synaptic dynamics and connectivity

The excitatory and inhibitory synaptic conductances induced by a presynaptic spike at time *t* = 0 are given by a bi-exponential function,

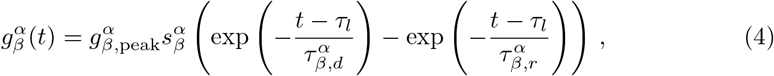

for *t* ≥ *τ_l_*, where *τ_l_* is the transmission delay, and 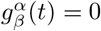 otherwise. 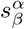 is a constant ensuring that 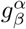 has its maximum at 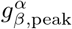. Suppressing the indices *α* and *β* we have

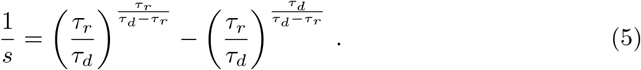

This setup is similar to that used in [44] and [25]. We show time courses for post synaptic potentials, conductances and post-synaptic currents in Fig 11, in the SI section Synaptic dynamics in CA1, for a fixed holding potential of −55 mV.

The two neuronal populations are coupled uniformly at random, with probabilities *p_αβ_*, where *α* ∈ {E, I} denotes the presynaptic population and *β* ∈ {E, I} denotes the postsynaptic population. If not mentioned otherwise, we set *N_E_* = 12000 and *N_I_* = 200, which approximately agrees with the neuron numbers in a typical CA1 slice, of thickness 0.4 mm and a volume of 0.057 mm^3^ [25]. The numbers respect the ratio of pyramidal cells to PV+ basket cells in rat CA1, which is approximately 60:1 [57]. To compute the connection probabilities *p_αβ_*, we assume that all connections between E and I cells are realized in the subset of CA1 neurons we consider. This means that we first determine how many inputs each E or I cell receives from the other cells in the population and then compute the connection probabilities using the numbers for *N_E_* and *N_I_* given above.

A hallmark of CA1 is its very sparse recurrent excitatory connectivity [56, 57]. We set *p_EE_* = 1.64% [56] so that for *N_E_* = 12000, every pyramidal cell receives input from approximately 200 other pyramidal cells.

For the I-to-I connectivity, we set *p_II_* = 0.2 [57, 79, 80], which, for *N_I_* = 200, means that each basket cell on average receives 40 synapses from other PV+BCs in the network. This value is obtained as follows: a single PV+BC contacts on average 64 other PV+ cells [79], of which 60% are basket cells [57], so that a single PV+BC contacts 38 other PV+BCs on average [25]. This corresponds approximately to *p_II_* = 0.2.

For the I-to-E synapses, we obtain several estimates, (i) based on the experimentally observed number of PCs that are postsynaptic to a single PV+BC and (ii) based on the experimentally observed number of PV+BCs that are presynaptic to a single PC. We first consider ref. [79]: (i) The conducted experiments in CA1 found that basket cells innervate between *D* = 1500 and *D* = 2000 pyramidal cells each. For *D* = 1500 we obtain, using the assumption of homogeneous and random connectivity, 1500 = *p_IE_N_E_* and thus *p_IE_* = 0.125. Analogously, *D* = 2000 yields *p_IE_* = 0.167. (ii) Ref. [79] further showed that 30 – 40 PV+ cells, of which 60% are PV+BCs [57], converge on a single CA1 PC. Thus, a single PC should be contacted by 18 ≤ *C* ≤ 24 PV+BCs. Under the assumption of homogeneous random connectivity, the mean number of PV+BCs contacting a single PC is given by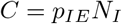, which, with the values for *p_IE_* obtained using estimate (i) above (0.125 ≤ *p_IE_* ≤ 0.167), would result in 25 ≤ *C* ≤ 34. This is slightly too large in comparison to the experimental values from estimate (ii). Therefore, a lower value of *p_IE_* = 0.1 seems also plausible. The meta-study [57] allows to obtain similar estimates: (i) It was shown that each PV+BC contacts on average 943 PCs. This fixes *p_IE_* because 943 =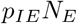, yielding *p_IE_* = 0.079. (ii) Ref. [57] further found that each pyramidal cell is innervated by 17 PV+BCs, such that 17 =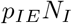, resulting in a value of *p_IE_* = 0.085, which is also close to 0.1. We thus fix *p_IE_* = 0.1, and consider values in the range from 0.1 to 0.2 as biologically plausible.

We first determine *p_EI_* for the E-to-I connectivity in an approach analogous to (i) above. We use the experimental result that each CA1 PC innervates 91 interneurons, of which approximately 14% are PV+BCs [57]. Therefore each PC diverges to innervate approximately *D* = 13 PV+BCs (*D* is chosen in analogy to estimate (i) for the I-to-E connectivity above). Thus, 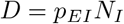, so that *p_EI_* = 6.5%. When trying to obtain the E-to-I connectivity in analogy to the approach (ii) above, one first notes that the number of excitatory synapses on a PV+BC is not known [57]. It has, however, been estimated that a hypothetical ‘average interneuron’ in CA1 receives 2211 excitatory boutons from local collaterals (i.e. local CA1 PCs) [57]. If each CA1 PC makes on average 3 synapses onto each postsynaptic interneuron [57], this means that each interneuron is contacted by approximately *C* ≈ 740 CA1 PCs. This results in 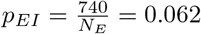. If each CA1 PC made only two synapses onto each postsynaptic interneuron, this would result in *C* ≈ 1106 and thus 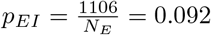. We therefore consider values 6.5% ≤ *p_E_* ≤ 10% as biologically plausible. For concreteness, we fix *p_EI_* = 0.1 in this study. This is at the upper end of the biologically plausible values, but acceptable because we will systematically vary the excitatory drive to the E population in our simulations: Decreasing *p_EI_* is similar to decreasing the number of active E cells since the latter reduces the number of the realized E-to-I connections that will have a postsynaptic effect. A decrease in the number of active E cells results in our models from changing the number *n_E_* of PCs that receive CA3 drive in model 1 and 2 and by changing the strength of the CA3 drive in model 3, see next section.

In summary, using the values available in the experimental literature [57, 79, 81–84] we obtained for our networks *p_EI_* = 0.1, so that each PV+BC receives on average input from 1200 PCs, and *p_IE_* = 0.1, so that each PC receives on average input from 20 PV+BCs. These connection probabilities are in good agreement with many previous computational studies of CA1 [36, 43, 44]. Notably, [25] use the same values for the connection probabilities except for a higher value of *p_IE_* = 0.3. Our more detailed choice for the connection probabilities differs from that in ref. [37], where a single value of *p* = 0.2 for all synapses was assumed. Finally, it differs from the all-to-all connectivity for all synapses except the E-to-E synapses assumed by [29].

To conclude, as a result of the considerations in this section we set *p_EE_* = 0.0164, *p_II_* = 0.2, *p_IE_* = 0.1 and *p_EI_* = 0.1 (if not mentioned otherwise).

### Excitation of pyramidal cells by CA3

Models 1 and 2 explicitly mimic sharp-wave input from CA3 without modeling CA3 [25]: a subset *n_E_* of the pyramidal cells is driven by time-dependent conductances *g*_ext_ with a Gaussian profile,

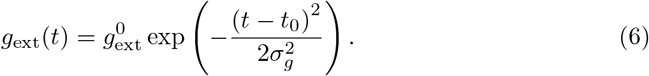

In model 3, CA3 is modeled as a population of *N*_*E*,CA3_ = 15000 excitatory neurons that each spike according to an inhomogeneous Poisson process with rate time course

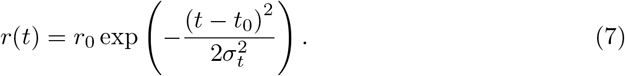

The connection probability between CA3 and CA1 is 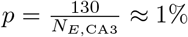 if not mentioned otherwise. This means that every CA1 pyramidal neuron on average receives input from 130 CA3 pyramidal neurons. The number is comparable to the typical number of inputs a CA1 neuron receives from CA3 during an SPW/R event (150 – 300 neurons, [85]). This is much lower than the number of CA3 neurons converging on a single CA1 pyramidal neuron (15000 – 30000, [57]). However, only approximately 1% of all CA3 pyramidal neurons are active during a SPW/R [59, 63]. Assuming that rat CA3 contains approximately 205000 PCs [57] and that the average connection probability from CA3 to CA1 (neglecting CA3 sublayer specificity) is between 1 and 8% [57], we obtain between 0.01 · 0.01 · 205000 ≈ 21 and 0.01 · 0.08 · 205000 = 164 inputs per CA1 PC, which is consistent with our choice for *p* given above. In light of this estimate, we consider one hundred to a few hundreds, but not thousand, inputs to a CA1 PC from CA3 to be realistic estimates. Our results do not depend crucially on the exact size of *p*. For example, a modest increase in *p* can be compensated for by a decrease in the average strengths of the peak dendritic currents (given by Eq 2) or the peak rate *r*_0_ in eq. 7.

As already mentioned above when introducing Eq 1, the impact of temporally coincident spikes from CA3 is nonlinearly amplified, in all *N_E_* excitatory cells: if in one of these cells 5 or more spikes arrive in an interval of *w_D_* = 2 ms [65], a dendritic spike is triggered. Irrespective of the coincidence detection mechanism, each CA3 spike impinging on a CA1 PC also causes a small depolarization (rise/decay time 1*/*2 ms, peak conductance 0.75 nA). The dendritic spike is incorporated in the model by the current *I*_dendritic_(*t*) that it generates in the soma (see Fig 2). We assume that the impact of the dendritic spike is stereotypical, i.e. 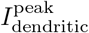 (cf. Eq 2) is for a given neuron constant over time and independent of the CA3 inputs that triggered it. This is plausible since dendritic spikes are stereotypical and couplings between dendrites and soma are reliable [71]. There are thus two sources of heterogeneity in the excitatory connections from CA3 to CA1. The first source is that the number of CA3 inputs impinging on a given CA1 cell is variable as only the connection probability *p* (see above) is fixed. The second source is the variable peak dendritic current 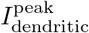 (cf. Eq 2) which is drawn, for each CA1 PC independently, from a lognormal distribution (Eq 2) with fixed parameters *μ* and *σ*. In our model, not every dendritic spike causes a somatic spike or even has a discernible influence on the soma (see Fig 2), which is in agreement with experimental observations [86]. Dendritic spikes have been observed in the apical dendrites of CA1 PCs during SPW/R [86]. These *large-amplitude fast spikes* [86] are initially fast and then broad, so that they are likely composed of both a sodium-dependent and a calcium-dependent component, which is generally less sensitive to synchronous coincident spiking [87], see also [88] for an example of a dendritic spike composed both of a calcium and sodium-dependent component. This is a caveat, because it might be that the large-amplitude dendritic spikes in [86] are less sensitive to synchrony in the input from CA3 or not caused by CA3 input at all, but by other mechanisms such as backpropagating action potentials [89]. Assuming that spikes are generated in the apical dendrites of CA1 pyramidal neurons during SPW/R by a coincidence-based mechanism, we can estimate the number of inputs *n_D_* required for its generation. To this end, we first define the dendritic coincidence rate as the number of inputs *n_D_* that have to arrive in the time window *w_D_* to cause a dendritic spike. To generate a dendritic spike, the mean CA3 afferent rate, which is given by *r*_0_*pN*_*E*,CA3_, should be at least equal to this event rate, thus we have the equality

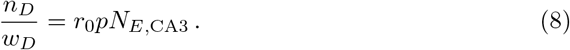

With *w_D_* = 2 ms, *pN*_*E*,CA3_ = 130 and 8 Hz ≤ *r*_0_ ≤ 10 Hz, this gives

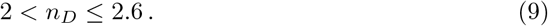

For *w_D_* = 3 ms, the estimate becomes 3 < *n_D_* ≤ 3.9. If additionally, *pN*_*E*,CA3_ = 150, we have 3.6 ≤ *n_D_* ≤ 4.5. This is lower than *n_D_* = 5 that we have assumed above, but on the same order of magnitude. This number of inputs *n_D_* and the time window *w_D_* are in agreement with the conditions for the generation of spikes in apical dendrites a realistic model of CA1 PCs [90]. Hence, in our model, few, but strong, CA3-CA1 synapses cause dendritic spiking. Our choice of parameters for the CA3-CA1 connectivity and the CA3 peak rate also entail that only neurons that receive more than average (*r*_0_*pN*_*E*,CA3_) CA3 input spikes will generate dendritic spikes. If not mentioned otherwise, there is no cutoff in the distribution of the peak dendritic currents. If a cutoff is imposed, after drawing the peak dendritic current for each neuron, all peak dendritic currents above the cutoff are set to 0 nA. We show a schematic summary of our three models in Fig 1. All simulations were performed with custom scripts in `brian2` [91].

## Results

In the following, we study the three minimal CA1 two-population models Fig 1 for the generation of high-frequency oscillations (HFOs) in the ripple range. The first quantities of interest are the frequencies of the network activity of the *I* and *E* population. We compute their means *f_α_* (*α* ∈ {*E, I*}) and standard deviations std(*f_α_*) (*α* ∈ {*E, I*}) across at least 10 network simulations with independent random realizations of external noise and connectivity. A small standard deviation indicates that the oscillation generated by the network is ‘robust’. This is a desirable property because it indicates that the oscillations are hardly affected by the ‘frozen’ noise caused by the random connectivity and the noise on the membrane voltage (cf. Eq 1 and Eq 3). In addition, to assess the firing activity of individual cells, we compute the mean number of spikes per active neuron *C_α_* and the fraction of active cells in the population *q_α_* (*α* ∈ {*E, I*}). This quantity is defined as the ratio of the number of cells which spiked at least once during the whole simulation and the total number of neurons *N_α_*, *α E, I*. We chose these basic statistical measures because they allow for an easy comparison with experimental values. In particular, computing spike counts instead of rates is suitable for our short simulations.

Following [22], where it was shown that exciting a small number of CA1 pyramidal cells is necessary and (in the presence of intact inhibition) also sufficient to induce a ripple oscillation, we first do not include any external excitatory drive to the inhibitory population of PV+BCs. That inhibition is also necessary [22] suggests a model of rhythmogenesis where the excited pyramidal cell subpopulation excites inhibitory cells, which then inhibit their excitatory targets. In such a model, the network frequency will depend on the recurrent E-to-I and I-to-E connections [37].

### Temporally homogeneous excitation of pyramidal cells (model 1)

We first study a model in which a subset of the PC population receives input that peaks at the same time *t*_0_, but has a different amplitude for each cell. The setup is similar to the indirect drive condition in [25]. In a minimal variant of the network, there is no external excitation of the PV+BC population, that is, the PV+BCs would be silent without input from the PC population. A subset of *n_E_* PCs is each excited by a time-dependent conductance (Eq 6), which peaks at *t*_0_ = 50 ms. The amplitudes 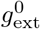 of the conductances are distributed according to a Gaussian with mean 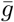and standard deviation 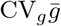 where CV*_g_* is the coefficient of variation, which is kept constant when we vary 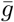.

To gain intuition, in Fig 3, we show network activities and histograms of spike counts for the two populations for a fixed level of excitatory drive and a fixed number of excited E cells 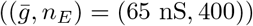. We see that E cells that are active tend to be active on multiple ripple waves (i.e. ripple cycles). This is because high external input amplitudes remain high over multiple ripple waves. Active E cells spike considerably more often than once or twice (on average they spike approximately 6 times in the event of Fig 3, see also Fig 4B for the corresponding average over events *C_E_*). I cells basically spike on every ripple wave (on average 6 – 7 times in Fig 3). In particular, nearly all I cells are active, *q_I_* ≈ 1. Most E cells, in contrast, remain silent, as evidenced by the large peak at 0 in the histogram for the E cell spike counts in Fig. 3 B, left panel.

**Fig 3.**
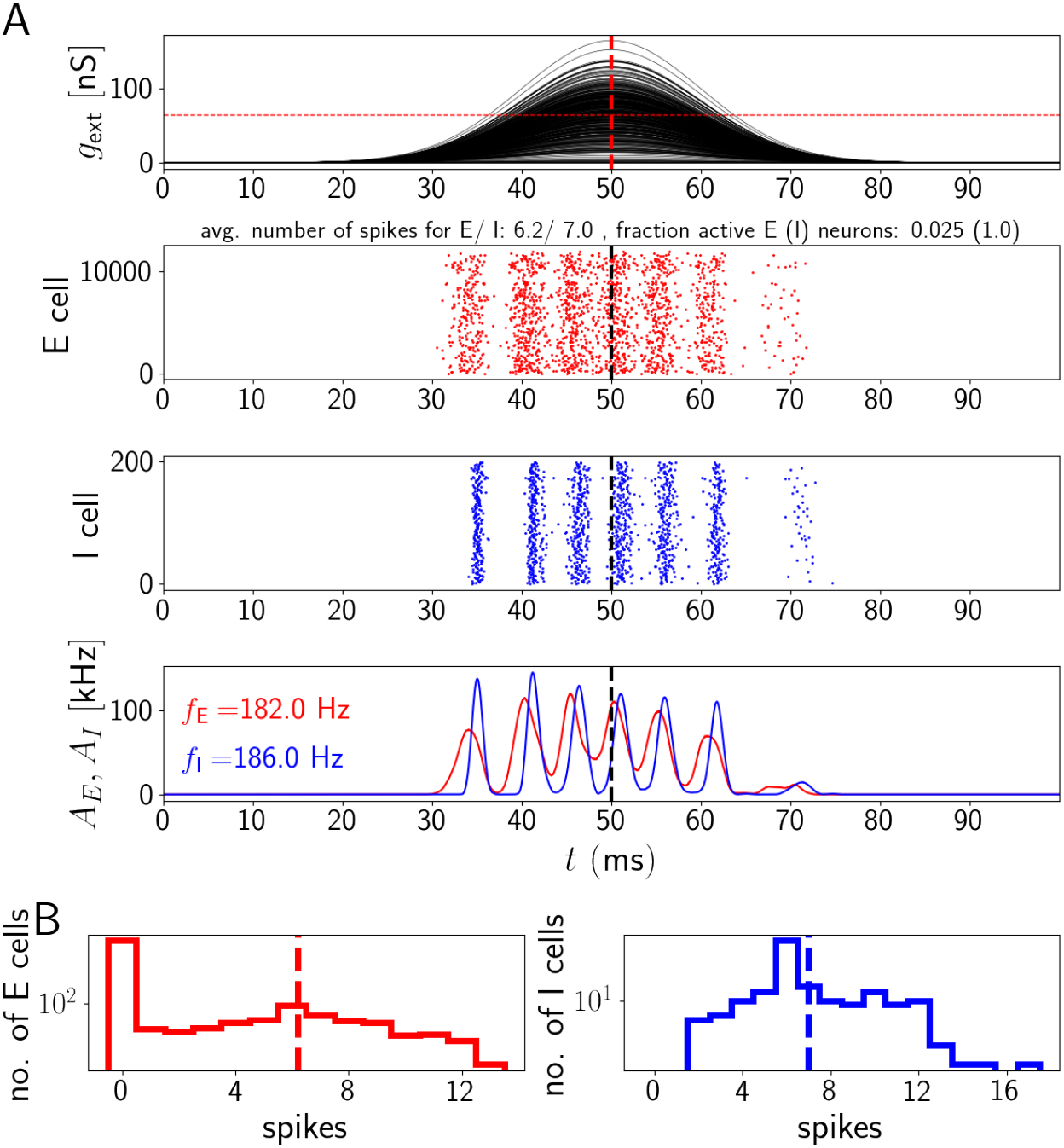
Network activity for temporally homogeneous excitation of E cells. The population activity of E and I cells oscillates at ripple frequency, but the active E cells contribute many spikes to a SPW/R event, like I cells. A, upper subpanel: Time courses of sharp wave input to the CA1 E cells (input conductances). The horizontal red dashed line indicates the mean 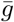 of the sharp wave amplitudes. Middle: spike rastergrams of the CA1 E and I cells (red and blue). Lower: population rates (red: population of E cells, blue: population of I cells). *f_E_* and *f_I_* in the lower panel give the oscillation frequency of the E and I population (spectrogram peak). B: Histograms of spike counts for E (left) and I (right) cells on a logarithmic scale. Parameter values as in Fig. 4, 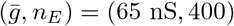.

**Fig 4.**
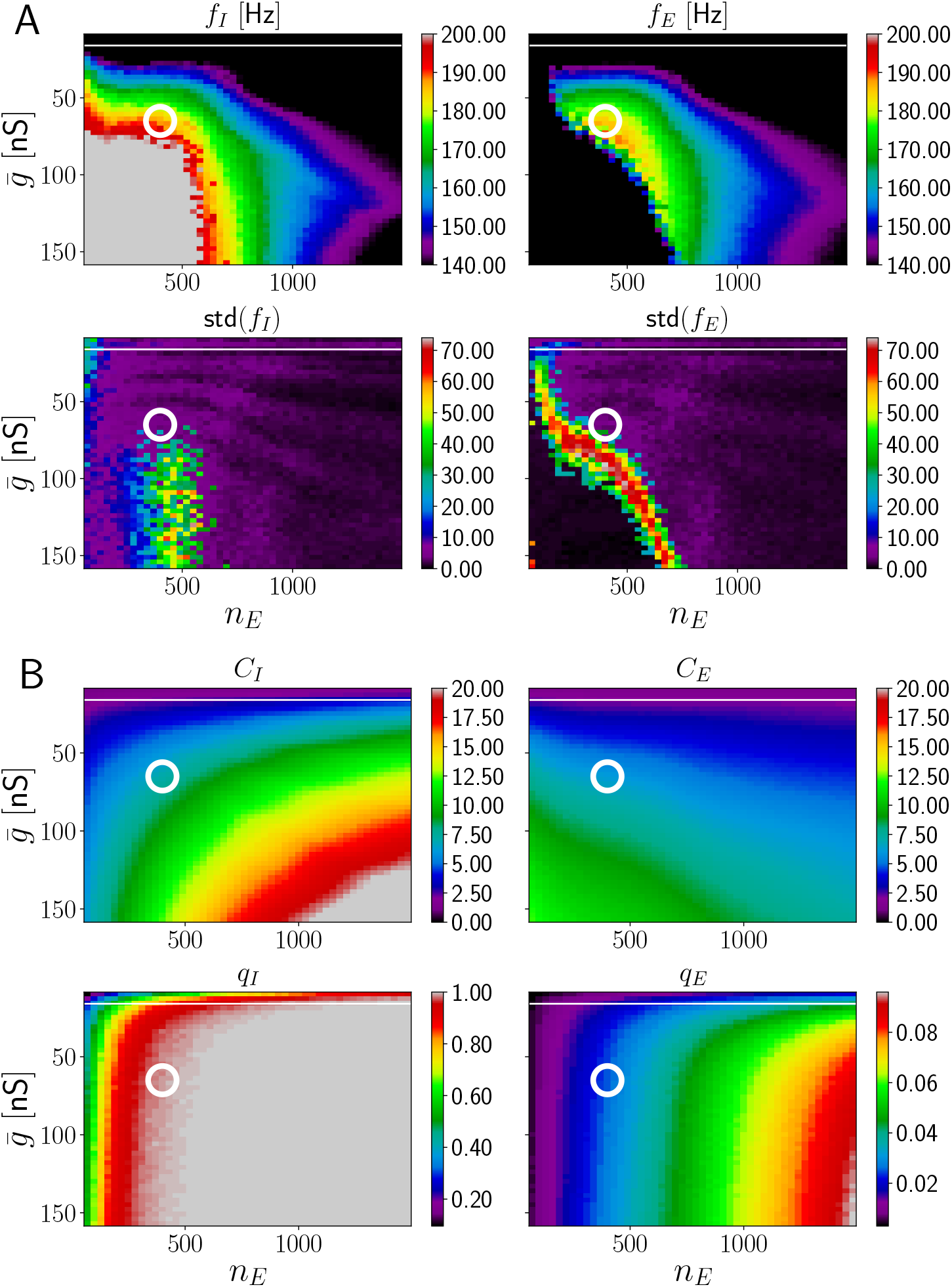
HFOs in networks with temporally homogeneous excitation of E cells. There is no overlap of the parameter region with reliable ripple oscillation generation and sparse spiking of E neurons. A, upper subpanel: Frequency of I and E population activity oscillations *f_I_*, *f_E_* as a function of mean drive 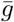 and number of excited E cells *n_E_* (average taken across different network realizations). Lower: Corresponding standard deviations across network realizations, std(*f_I_*) and std(*f_E_*). B, upper subpanels: mean number of spikes per active inhibitory (excitatory) neuron *C_I_* (*C_E_*). Lower: fraction of active neurons (*q_I_* and *q_E_*). The range for *f_I_* and *f_E_* displayed in detail by different colors is the ripple range, [140, 200] Hz. The range for both *C_I_* and *C_E_* is [0, 20]. Frequency values below (above) this range is indicated in black (grey). To guide the eye, the white horizontal line indicates a border of the range of biologically plausible *C_E_*; this range lies completely above the line, which is at 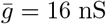. The white circle indicates 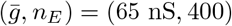, the parameters used in Fig 3. Standard deviation of the sharp wave peak conductance is CV*_g_* = 0.5, width of sharp waves *σ_g_* = 10 ms.

To confirm that this is a generic problem of model 1 and not due to our choice of 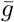 and *n_E_*, we systematically varied these parameters. The results are shown in Fig. 4. The rainbow coloring in Fig 4 A, upper subpanels, marks the region where the expected PC and PV+BC population activity oscillates in the ripple frequency range (upper panels in A). The magenta part of the lower subpanels indicates the region where the standard deviation of the oscillation frequencies across different network realizations is small (lower panels in A). Networks with parameters that fall into both regions can be expected to generate robust ripple frequency oscillations. However, in the entire region where the expected oscillation frequency is in the ripple frequency range, an active pyramidal cell spikes on average more than 5 times during the whole ripple cycle (Fig. 4 B, Fig 3 B). This is in marked disagreement with experimental findings [20], which show that most pyramidal cells fire once or twice during the ripple cycle. More recent experiments even observed that E cells typically spike only once per ripple [22]. The white horizontal line in Fig 4A roughly delimits the region where E cells spike sparsely. Anywhere below this white line, i.e. for higher values of 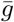 *C_E_* is larger than two, which is biologically implausible.

Thus, in our model 1, in the absence of external feedforward excitatory drive to inhibitory cells, it is not possible to reach the ripple frequency range with sparse firing of pyramidal cells by changing 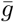 and *n_E_*. This remains true if *p_IE_* is increased from 0.1 to 0.2 (Fig. 12, SI, section Higher I-to-E, I-to-I connectivity and broader sharp waves in model 1). We increased *p_IE_* because one might expect that more inhibitory inputs received by each PC result in fewer E spikes; however, there still is no region where HFOs in the ripple range co-occur with small values for *C_E_*. Increasing *p_IE_* further and also increasing the width of the sharp wave *σ_g_* to distribute the firing of the PCs does not result in lower firing of the E cells but instead lowers the frequency of the network oscillations to a maximum of approximately 170 Hz (Fig 13, SI, section Higher I-to-E, I-to-I connectivity and broader sharp waves in model 1). We note that larger network frequencies can always be obtained by increasing *p_II_* and decreasing *p_IE_* (Fig 14, SI, section Higher I-to-E, I-to-I connectivity and broader sharp waves in model 1); however, there still is no region where HFOs in the ripple range co-occur with small values for *C_E_*.

Finally, we include a strong feedforward excitatory drive to inhibitory cells. Such strong drive to the I cells renders our model an IR model (see Introduction). This is because the I cell population already generates the observed oscillation, while the impact of the E cells is small. This is shown in Fig 15 (SI, section Strong feedforward excitation to I cells in model 1), where each PV+BC is driven with a conductance as in Eq 6 with *σ_g_* = 10 ms. All I cells get the same amplitude 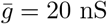 which results in robust oscillations at *f_I_* ≈ 165 Hz in the absence of the pyramidal cell population. With this modification, we now observe a region in parameter space where, for small *n_E_* and 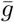 robust HFOs in the ripple range with small *C_E_* < 2 are reached. Analogous to Fig 3, we show representative network activity in Fig 16 (SI, section Strong feedforward excitation to I cells in model 1). Due to the weak drive and strong inhibition, E cell spiking is very sparse: in Fig 16, less than 30 E cells are active, which is less than 1% of the whole population. Therefore, E activity does not discernibly influence the rhythmic I spiking. I cells start spiking before E cells on every ripple cycle. Although E cell activity is sparse, some E cells still spike more than twice during the ripple (Fig 16 B, SI, section Strong feedforward excitation to I cells in model 1). Thus, in the presence of strong feedforward excitation to the inhibitory cells, model 1 leads to both network and single cell statistics consistent with the experimental data. However, the experiments of [22], find that strong drive to the I cells is insufficient to generate HFOs in the ripple range *in vivo* and in fact, abolishes ongoing ripples (Fig 5E in [22]). Moreover, with the inclusion of a strong feedforward drive to the I cells, I cells start to spike before E cells in each ripple, which is not observed in experiments [20, 22]. Hence, we do not consider strong excitatory feedforward drive to the I cell population as a biologically plausible option to decrease the activity of E cells in our model 1. Whether such strong feedforward drive during SPW states is realized in *in vivo* remains, however, an open question [92].

**Fig 5.**
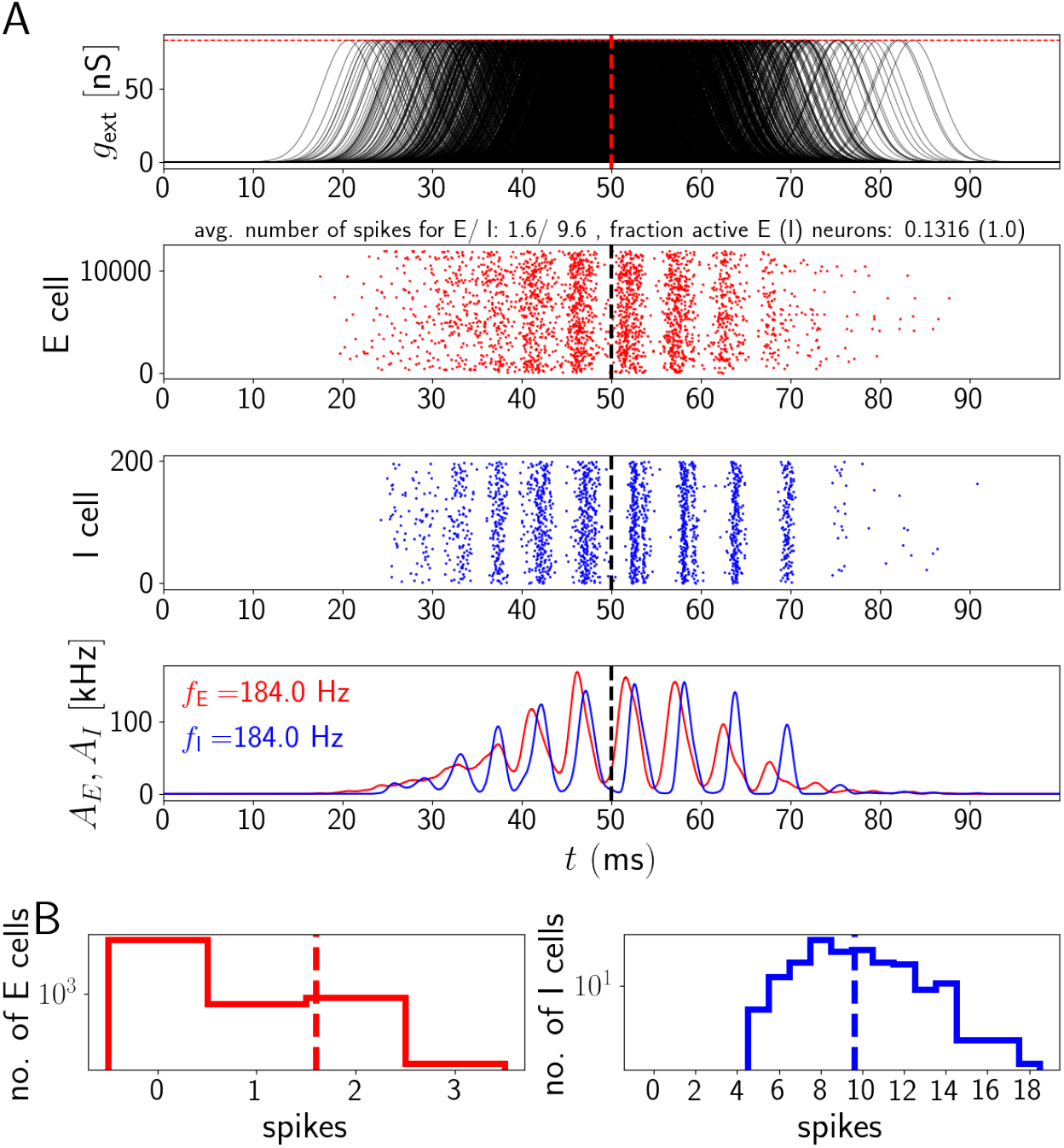
Network activity for temporally inhomogeneous excitation of E cells. Each E cell receives an input pulse whose duration is comparable to the interval between two ripple waves. The peak times of the pulses are distributed. E and I populations spike at ripple frequency, active E cells typically contribute one to two spikes to an event. A: Input pulses, spike rastergrams and network activities, displayed as in Fig 3. B: Histograms of spike counts during the displayed event. Parameter values as in Fig 6, 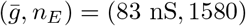.

In conclusion, model 1 does not seem suitable to describe SPW/Rs, where high network frequencies in the range from 140 to 200 Hz should co-occur with sparse firing of PCs.

### Temporally inhomogeneous excitation of pyramidal cells (model 2)

We next ask whether the biologically implausible frequent E cell firing during ripple frequency oscillations in our two-population models can be avoided by providing excitation to CA1 pyramidal cells such that the input pulses are shorter and received at different time. For this we distribute the input peak times across the pyramidal neurons according to a Gaussian distribution, 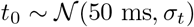. The individual excitation pulses have each a width of *σ_g_*. For simplicity, we assume that all pulses have the same amplitude, 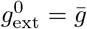 such that all PCs receive the same amount of excitation, but at different points in time.

A sample output of this model for *σ_g_* = 3 ms is shown in Fig 5. Robust oscillations in the ripple range are reached in the region around 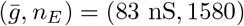. In contrast to Fig 4, active E cells typically contribute 1 or 2 spikes to an event.

We confirm in Fig 6 that there is a broad region in the parameter space spanned by 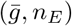 where HFOs in the ripple range co-occur with sparse firing of E cells. This region lies at intermediate values of 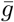. An increase (decrease) in 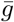 has to be accompanied by an increase (decrease) in *n_E_* to stay in this region. Thus, higher levels of excitation have to be distributed over more E cells to stay in the ripple range.

**Fig 6.**
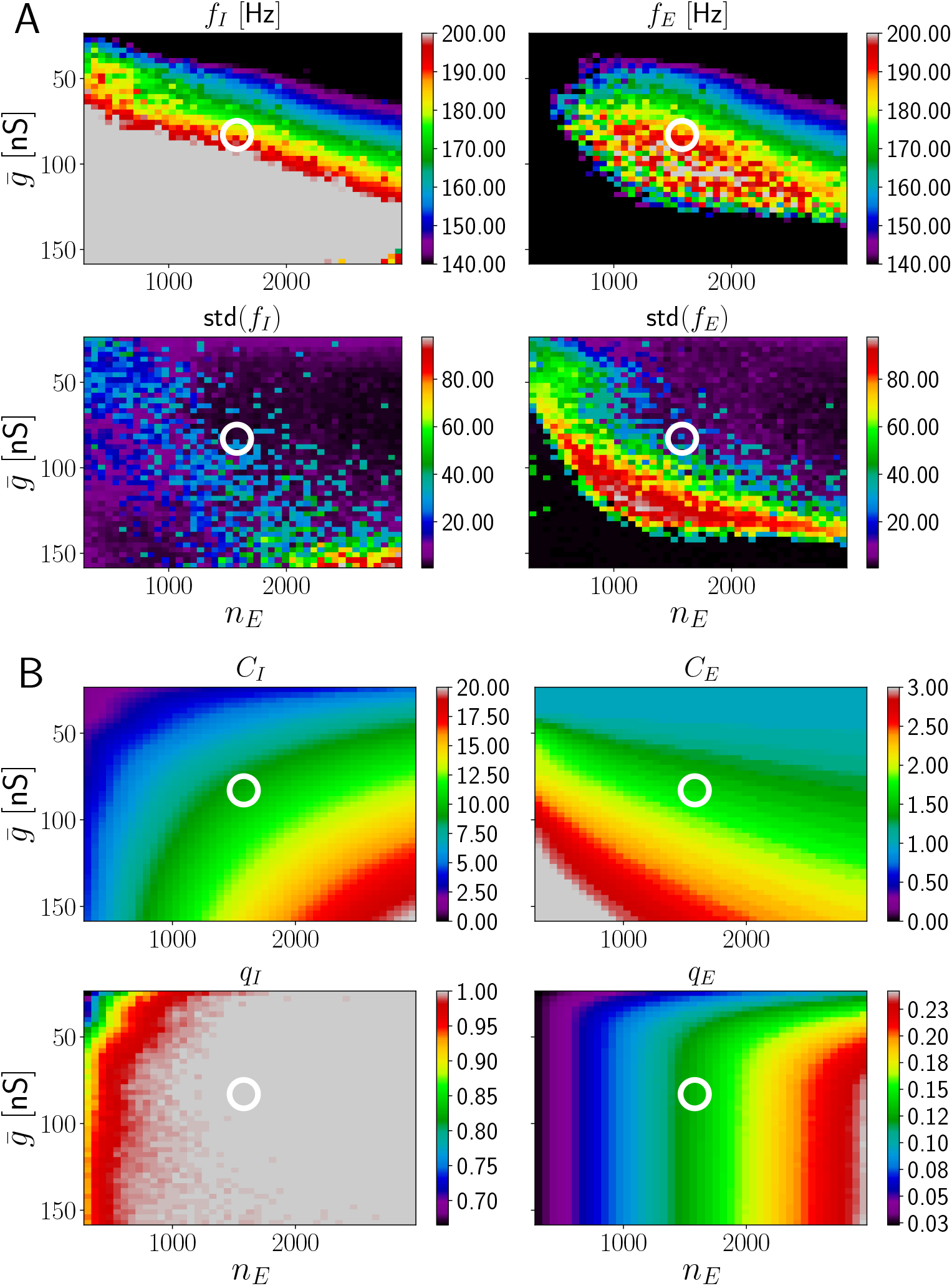
HFOs in networks with temporally inhomogeneous excitation of E cells. Ripple oscillation generation with sparse spiking of E neurons is robust in networks where the E cells are driven by short input pulses. Figure layout as in Fig 4. A: Network frequencies and standard deviations. B: Number of spikes per active neuron and fraction of active neurons. The ranges for *C_I_* and *C_E_* are [0, 20] and [0, 3], respectively. The white circle is located at 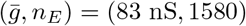 Parameters: *σ_t_* = 10 ms, *σ_g_* = 3 ms.

To maintain sparse firing of E cells, individual external input pulses need to be narrow. In Fig 17 (SI, section Broader pulses in model 2), we increase their width to *σ_g_* = 5 ms, which results in more E cell spikes per active E cell (between 2 and 3 spikes on average in the region where ripple frequency oscillations are generated).

As shown before for model 1, higher oscillation frequencies can be reached by increasing *p_II_*. Additionally increasing *p_IE_* renders the oscillations more robust and the region in parameter space where HFOs in the ripple range are generated increases (Fig 18, SI, section Larger I-to-E, I-to-I connectivity and broader spread of individual input pulses in model 2). We show in Fig 19 (SI, section Larger I-to-E, I-to-I connectivity and broader spread of individual input pulses in model 2) that the network dynamics and statistics stay realistic when we, in addition to *p_II_* and *p_IE_*, also increase *σ_t_*, which controls how broadly the pulse peaks are distributed over time (Eq 6). In Fig 20 (SI, section Larger I-to-E, I-to-I connectivity and broader spread of individual input pulses in model 2), we show the network dynamics and statistics when we decrease *p_II_* back to 0.2 (keeping *p_IE_* = 0.2) compared to Fig 19. Again, there is a regime where high network frequencies and sparse firing of pyramidal neurons coexist.

In conclusion, for model 2 the generation of HFOs in the ripple range with sparse firing of pyramidal cells does not depend much on the details of how the recurrent I-to-E and E-to-I loop are set up (within the limits of biological plausibility). Instead, the main determinant for the sparseness of pyramidal cell activity are the properties of the afferent CA3 drive. In the next section, we show how dendritic spikes present in a strong basal or apical dendrite of CA1 pyramidal cells might give rise to temporally localized input that has a strong influence on somatic spiking output.

### Dendritic spikes provide short windows of excitation for pyramidal cells (model 3)

Dendrites of CA1 pyramidal cells can, when excited with sufficiently synchronous and spatially clustered inputs, generate dendritic spikes that propagate to the soma and may cause an action potential there [65–67, 93]. Dendritic spikes are of particular interest when modeling ripple oscillations in CA1 because both apical and basal dendrites of CA1 pyramidal cells are targeted by the Schaffer collateral excitatory input from CA3 [1, 94]. Moreover, it was shown that *in vivo*, dendritic spikes occur during sharp-wave activity [86]. Ref. [67] suggested that dendritic spikes observed in the apical dendrites of CA1 pyramidal cells endow the cell with unique information processing capabilities during SPW/Rs. Specifically it was shown that these dendritic spikes result in precise somatic spikes with low temporal jitter. This lead to the conjecture that dendritic compartments receiving input clustered both in space and time perform supra-linear dendritic integration during SPW/R events. The dendrites might then act as feature detectors on the CA3 input and determine the neuronal action potential output. This would be largely independent of the mean input strength in contrast to the output during theta oscillations [67].

Consistent with these findings and suggestions, we propose that dendritic spikes may provide the short-term excitation that we found to be necessary for HFOs with sparse E cell firing in model 2 (see Fig 6 and Fig 17). For this, we model the generation of dendritic spikes in the E neurons that receive input from CA3. Since dendritic spikes are generated by coincident spike arrivals, we introduce a simple model of CA3 with *N*_*E*,CA3_ pyramidal cells, each of which fires according to an inhomogeneous Poisson process. This has a peaked rate profile with the same amplitude and width across all CA3 neurons (see Eq 7, the peak is at *t*_0_ = 50 ms, width is *σ_t_* = 10 ms). When there are at least five input spikes arriving within a time window of 2ms, a dendritic spike is elicited. Dendritic spikes impact a neuron’s somatic voltage in a stereotypical fashion. Between neurons, their impact varies: the peak current elicited by a dendritic spike in the soma is distributed according to a lognormal distribution (Eq 2) across the CA1 E cells. Each CA1 PC receives inputs from a small fraction of all CA3 pyramidal cells. We set the connection probability to 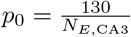, so that the average number of CA3 inputs to a CA1 E cell is 130. The size of the modeled CA3 population is *N*_*E*,CA3_ = 15000 and their peak rate is *r*_0_ = 8 Hz [64, 95, 96] (see Materials and Methods for further details).

Besides their contribution to dendritic spikes, the inputs from CA3 elicit short excitatory postysnaptic currents with an excitatory postsnaptic potential of 0.4 mV when the neuron is otherwise held at −55 mV. Compared to model 2, we have decreased the synaptic latency from I to E cells from 1.0 ms to 0.5 ms. This stabilizes the oscillations, but is not essential, see Fig 24 in the SI. That it is helpful can be intuitively explained as follows: The spike rate of the CA3 population (given by Eq 7) is towards its peak sufficiently high to generate widespread spiking in the CA1 pyramidal cells. Fast recurrent inhibition within CA1 is required to terminate a single pyramidal ripple wave; in the absence of inhibition it would go on as long as CA3 spiking is strong enough. Inhibition has to impinge early enough on the excitatory population on every ripple wave to prevent an excess of excitation and thus a slowing of the oscillation due to resulting excessive inhibitory spiking. Therefore, a parameter change that lets recurrent inhibition arrive earlier decreases the number of E spikes and renders the oscillations more pronounced and robust. Any mechanism that decreases the width of the excitatory ripple wave will have a similar effect. In addition to the PV+BCs considered here, such silencing of pyramidal cells could be provided by bistratified cells [97, 98] or feedforward inhibition [99].

Fig 7 shows a representative simulation of a model 3 network generating HFOs in the ripple range accompanied by sparse firing of pyramidal cells. Whereas the mean number of spikes in the active E cell population is between 1 and 2 (approximately 1.5 in Fig 7) and most active E cells spike once or twice, a small fraction of neurons spikes 5 times or more during the SPW/R event (left histogram in Fig 7 B). Such ‘bursting’ behavior is caused by large values for the peak of the dendritic current and/or exceptionally high synchrony in the presynaptic spike trains impinging on a CA1 PC. We will see below that it is not crucial for generating SPW/Rs in our models (SI, Figs 25, 26, 27 and 28). Bursting of pyramidal cells in CA1 was observed *in vivo* during slow wave sleep [60, 100], in particular during SPW/Rs [39, 101].

**Fig 7.**
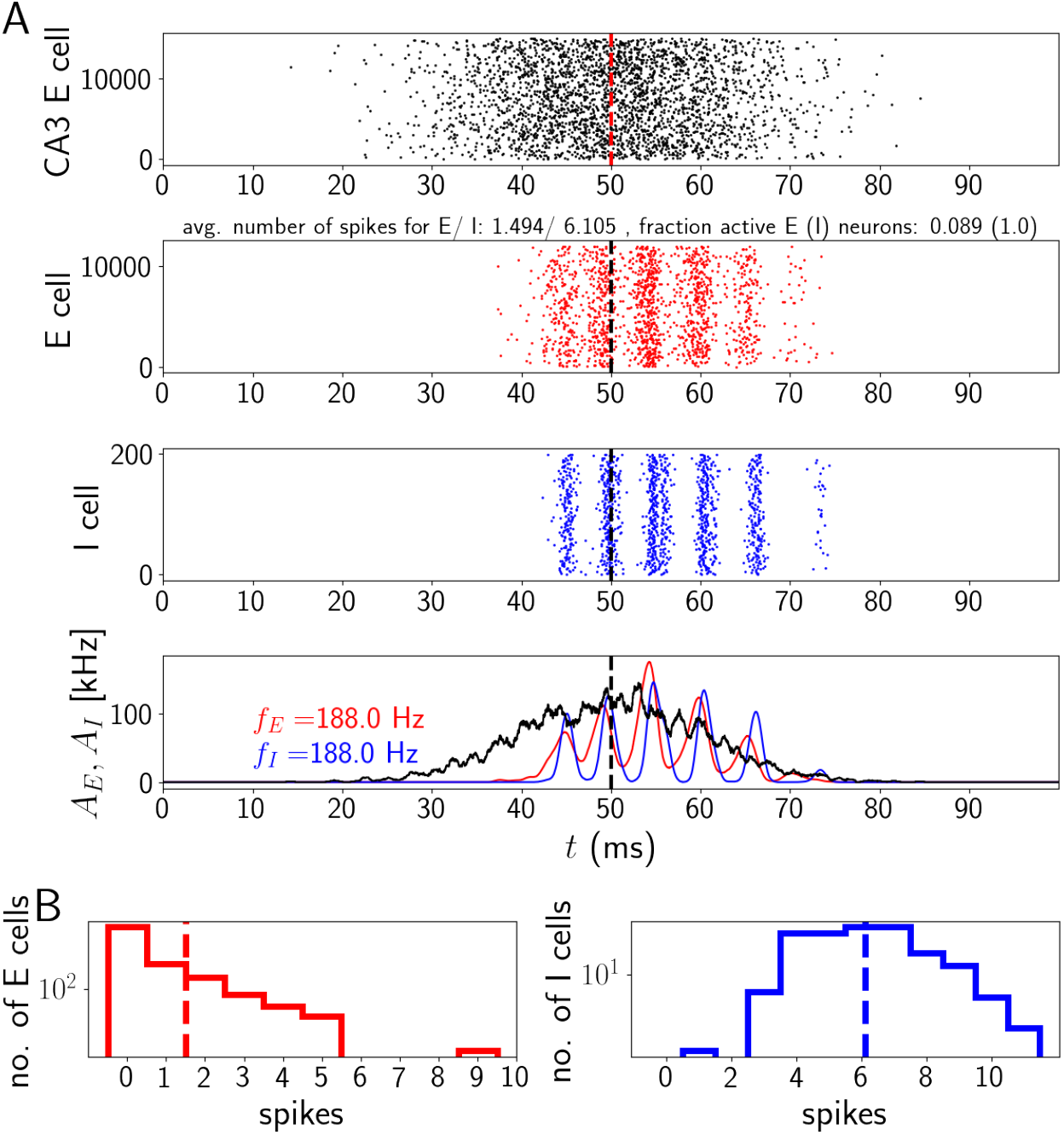
Activity in a network with nonlinear dendritic excitation of E cells. CA3 inputs can cause dendritic spikes in the E cells receiving them. E and I populations spike at ripple frequency, active E cells typically contribute one or two spikes to an event. A, upper subpanel: Spikes generated by CA3. Middle: spike rastergrams of the CA1 E and I cells (red and blue). Lower: population rates (red: population of E cells, blue: population of I cells, black: population of CA3 cells). B: Histograms of spike counts. Parameter values as in Fig 8, (*σ, μ*) = (0.75 nA, 0.0 nA). Fraction of active E/I cells (*q_E_, q_I_*): 0.089, 1. Number of spikes per spiking E/I cell (*C_E_, C_I_*): 6.1, 1.5.

Why does model 3 generate sparse E cell firing in contrast to model 1, if in both models the CA1 E cells receive a broad sharp wave input? In model 3, CA1 receives noisy Poisson spiking and each E neuron has a short integration window for dendritic spike generation. Such a short window generates little averaging over the input spikes. Therefore the fluctuations in the number of spikes arriving within the window are large compared to the average. When the temporal density of input spikes to a CA 1 E neuron is by chance exceptionally high, a dendritic spike is generated. This happens rarely, but not with negligible frequency. The resulting sparse dendritic spiking provides narrow, pulsatile, strong inputs to these E cells, which leads like in model 2 to their sparse spiking.

In Fig 8, we systematically vary the parameters *σ* and *μ* of the lognormal distribution for the dendritic spike impact strength (Eq 2). For intermediate values of both *σ* and *μ* (rainbow colored band left and right of white circle in Fig 8 A upper panels), the system robustly generates fast oscillations in the ripple range with realistic single-neuron firing statistics. To stay in the ripple range, an increase in *μ* can be compensated by a *decrease* in *σ*. For a fixed value of *μ*, an increase of *σ* beyond a certain value renders the oscillations non-robust: the oscillation frequency varies considerably across different network realization (Fig 8 A lower panels).

**Fig 8.**
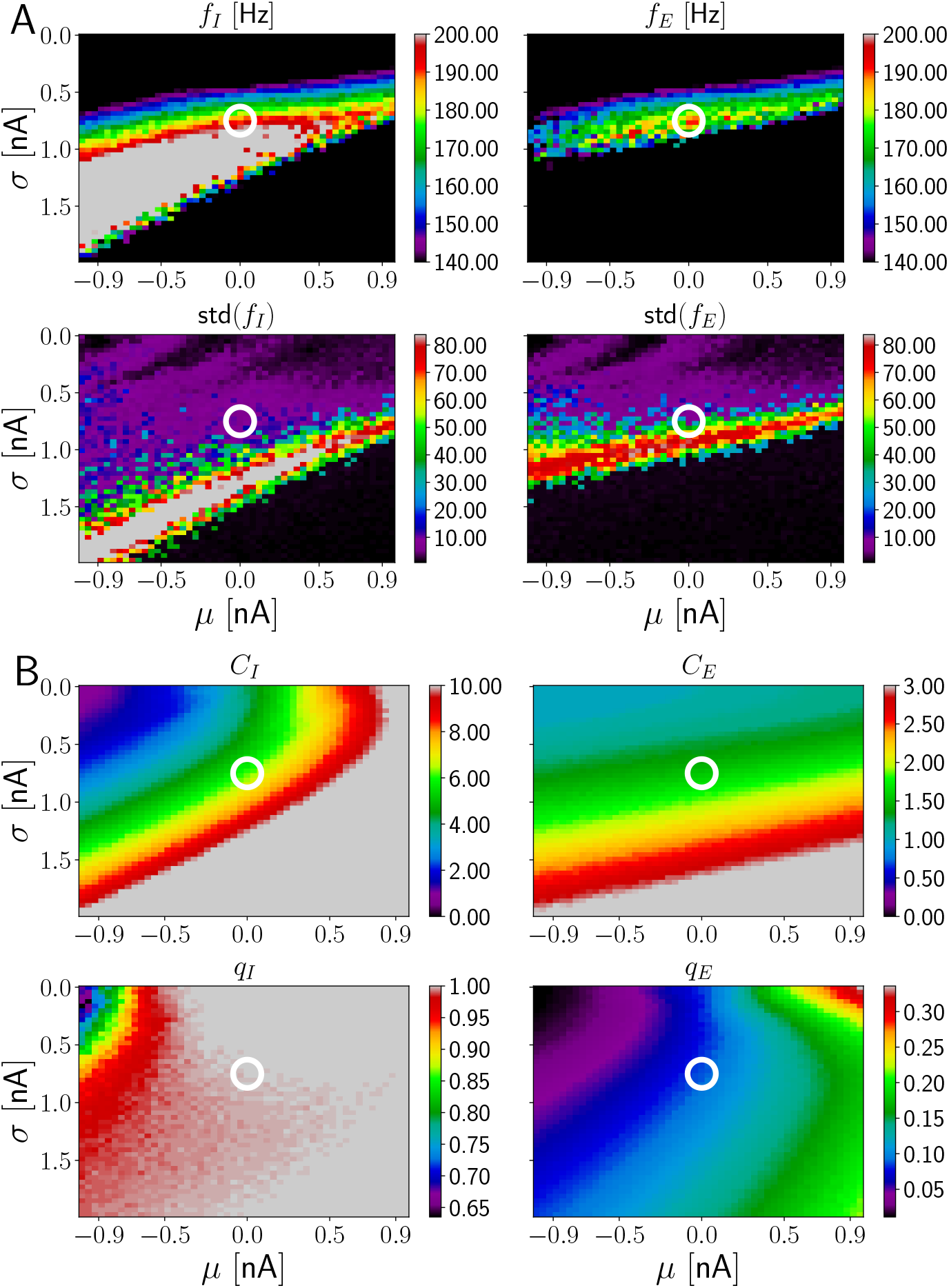
HFOs in networks incorporating dendritic excitation. Depending on the properties of the distribution for the peak dendritic current, a region with reliable ripple oscillation generation and sparse spiking of E neurons exists in model 3. The layout of the figure is similar to Fig 4; the changed variables are now the parameters of the distribution of dendritic spike impact strength (Eq 2). A: Network frequencies and standard deviations as a function of the parameters *μ* and *σ* of the lognormal distribution for the peak current induced in the soma by a dendritic spike (Eq 2). The range displayed in detail by different colors is the ripple range, [140, 200] Hz. Frequencies above and below are shown in gray and black. B: Number of spikes per active neuron and fraction of active neurons. The displayed ranges for *C_I_* and *C_E_* are [0, 10] and [0, 3], respectively, values above are colored in gray. The white circle is located at (*σ, μ*) = (0.75 nA, 0.0 nA), the parameter values of Fig 7.

E cell spiking is sparse all over the range where ripple frequency oscillations are generated: *C_E_* is approximately constant, between 1 and 2 (Fig 8 B). The amount of I cell firing depends on the precise values for the parameters of the peak dendritic current. For negative values of *μ* and *σ* ≈ 1 nA, I cells fire on average three times per ripple. Moving along the diagonal where ripple oscillations are generated to values of *σ* ≈ 0.5 nA and *μ* ≈ 0.9 nA, increases *C_I_* to more than 10 spikes per ripple. The range in between appears consistent with the experimental observation that CA1 PV+BCs typically spike on every ripple wave.

We now discuss the impact of certain selected parameter changes on model 3 to obtain an understanding of the robustness of our findings (SI, section Higher *p_IE_*, truncation of distribution for 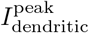 higher I-to-E synaptic latency and lower E-to-I peak conductance in model 3). Simulations for higher I-to-E connection probability (*p_IE_* = 0.3 instead of *p_IE_* = 0.1) are shown in the SI, Fig 21. The band where HFOs in the ripple range occur has moved to slightly larger values of *σ* compared to Fig 8. E cell firing is sparse within the band. Thus, model 3 is robust to a higher I-to-E connection probability. We argue in Materials and methods that *p_IE_* should be between 0.1 and 0.2 for the network sizes we consider in this paper. Given that HFOs in the ripple range are still observed with a higher value *p_IE_* = 0.3, we are confident that the results of model 3 do not hinge critically on the precise value of the I-to-E connection probability.

We next increase the I-to-E synaptic latency from 0.5 ms to 0.9 ms (SI, Fig 22). The maximal frequency reached by the E population is reduced to approximately 140 Hz and the region where this occurs is shifted to smaller values of *σ*. E cell firing remains sparse. This indicates that in model 3 HFOs in the ripple range (frequencies > 160 Hz) rely on fast synaptic transmission from the I to the E population.

We also decreased the peak E-to-I conductance 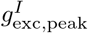 from 3 to 1 nS still in the experimentally observed range [83, 84]. This results in more robust oscillations in the ripple range (SI, Fig 23). Further, the region in the (*σ*, *μ*) parameter space where HFOs exist is enlarged compared to Fig 8, it now has the shape of a diagonal stripe extending from negative values for *μ* and large values for *σ* to positive values for *μ* and small values for *σ*. This appears broadly consistent with in *in vivo* experiments in transgenic mice, which found increased ripple power when reducing AMPA receptor-mediated excitation on PV+BCs [102].

Next, we asked how important the comparably small synaptic I-to-E latency (*τ_l_* = 0.5 ms in Fig 8 instead of 1.0 ms in models 1 and 2) is for the functioning of model 3. We find that an increase in *τ_l_* can be compensated by a decrease in the AMPA rise time on I cells (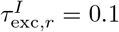 ms instead of 0.5 ms, SI, Fig 24). We note that additionally the peak E-to-I synaptic conductance was decreased (as in SI, Fig 23). The changes do not impact the ability of model 3 to generate HFOs in the ripple range with sparse firing of E cells. We conclude that model 3 does not critically hinge on the smaller value for the I-to-E synaptic latency used in Fig 8.

Finally, we study whether the large values at the tail of the lognormal distribution of dendritic spike impacts are important. We thus truncate the distribution given by Eq 2 such that values of 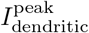 larger than 4 nA are mapped to 0. With this truncation, a single dendritic spike can generate at most, but barely, two somatic ones (see Fig 2 C for an example showing two somatic spikes as a response to one dendritic spike with a peak amplitude of 5 nA). Thus, with this truncation, there can be no somatic bursting due to a single dendritic spike. We find that this reduces the maximal frequency that can be reached (SI, section Higher *p_IE_*, truncation of distribution for 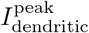 higher I-to-E synaptic latency and lower E-to-I peak conductance in model 3, Fig 25). However, frequencies around 170 Hz can still be reached and the standard deviation of the frequencies is low in that region, albeit higher than in Fig 8. Additionally, we found that the low ripple range (~ 150 – 160 Hz) can still be reached when the truncation is introduced at 3 nA, but not when it is introduced at 2 nA (not shown).

We additionally study a modified way to truncate the distribution for the peak dendritic current. Instead of setting values above the cutoff to zero, we re-sample values from the lognormal distribution until each value for the peak dendritic current is smaller than 4 nA. This lead to results similar to Fig 25 (SI), but with higher frequencies in the ripple range around 190 Hz (see red region next to white circle in Fig 26 A upper panels in the SI). The fact that ‘outliers’, which are eliminated by the truncation, are not crucial for the generation of HFOs in model 3 also suggests that our choice for the distribution of the peak dendritic currents (Eq 2) is not unique. Indeed, we find that results similar to Fig 8 can be obtained with a Gaussian distribution for the peak dendritic current that has the same mean and standard deviation as the corresponding lognormal distribution (Fig 27 in the SI). We also find that increasing the absolute refractory period of each CA1 PC to the unrealistically high value of 200 ms still results in networks generating robust HFOs in the ripple range (Fig 28 in the SI). With this absolute refractory period, the firing of the E cells is certainly sparse as they can at most spike once during a simulation of duration 100 ms. This proves that while there can be bursts of CA1 PC activity in our model, they are not necessary for the generation of HFOs in the ripple range.

In conclusion, high values for the peak dendritic current leading to somatic bursts (multiple PC spikes during one ripple wave) aid the model in generating fast and robust HFOs. They are, however, not necessary: after truncating the lognormal dendritic spike strength distribution, replacing it with a Gaussian one, or preventing E neurons altogether from generating multiple spikes by a long somatic refractory period, robust HFOs in the ripple range are still generated.

### Intrinsic CA1 replay by pulse and gap coding

A prominent feature of SPWRs in CA1 is that they occur in conjunction with replay of activity sequences from previous periods of exploration and learning [2, 101, 103–106]. Recent experimental results show that CA1 can generate sequence replay intrinsically without structured input from CA3 [39]. Further experimental evidence indicates that place cells in CA1 and CA3 have different properties [105, 107]. One possible explanation for the intrinsic replay in CA1 is that it is based on recurrent excitatory connections that are amplified due to dendritic spikes in the basal dendrites of CA1 [43, 44]. In this model, pulses of excitatory spikes travel along pathways in the sparse recurrent connectivity, which is enabled by amplification by dendritic spikes in the basal dendrites. Replay of spike sequences might also arise from continuous attractor dynamics. Here a localized bump of activity in the neural tissue moves around because of asymmetric synaptic connections, short-term plasticity, or adaptation mechanisms [108–112]. Sequential network structures might guide these bumps to replay certain sequences. Due to the very sparse excitatory recurrent connectivity, also propagation of inhibitory spikes pulses in an essentially inhibitory network as suggested for the striatum [113], might be considered biologically plausible for CA1. However, given the high firing rate of I neurons during ripples (as mentioned before, PC+BCs fire on nearly every ripple wave [1, 20]), it seems unlikely that their population activity forms sequential patterns similar to that of PCs. In support of this, it is also known that I cells have broader, more unspecific place fields than E cells [85, 99] (see however [114] and [115]).

Motivated by the high sparseness of excitatory-excitatory connectivity in CA1 and the supposed absence of replay in inhibitory activity, we propose an alternative model for how specific sequences may be stored and replayed. The networks are a modification of model 3, in particular there are dendritic spikes amplifying the input from CA3 to CA1. We expect that these are not needed, i.e. any form of strong feedforward excitation on CA1 PCs (for example the form used in model 1) should be sufficient for our replay model. Like the proposed ripple generation mechanism, sequence replay is based on the prominent excitatory-inhibitory and inhibitory-excitatory connectivity in the hippocampal region CA1. The basic idea is as follows: CA3 excites an initial group of CA1 E cells to spike within a short time interval [116]. This group excites all CA1 I cells involved in the dynamical pattern (‘pulse coding’), except for a group that would inhibit the second group of E cells. Due to the resulting disinhibition (‘gap coding’), this second group of E cells becomes active and excites all I cells except for a group, which would inhibit the third group of E cells. Therefore this group becomes active and so on (Fig 9 A). A generalization to continuous, non-grouped activity sequences [31, 44] seems straightforward. Further alternative concepts for replay, which we do not consider as they seem less biologically plausible for CA1, are presented in the SI. The first such concept (Fig 30 A) is a two-population model, which assumes that in addition to E cells also the I cells spike in a sequential fashion: the active group of E cells excites the next group of I cells, which inhibits all E cells except those that fire in the next step. The second concept is an inhibition-first model, which generates sequences by pure gap coding in the I population (Fig 30 B): groups of I cells become sequentially inactive, since they receive at some point more inhibition than their peers because of structured inhibitory connectivity. The resulting disinhibition of E cells leads to their sequential activity, but their spiking does not contribute to maintain the sequence.

**Fig 9.**
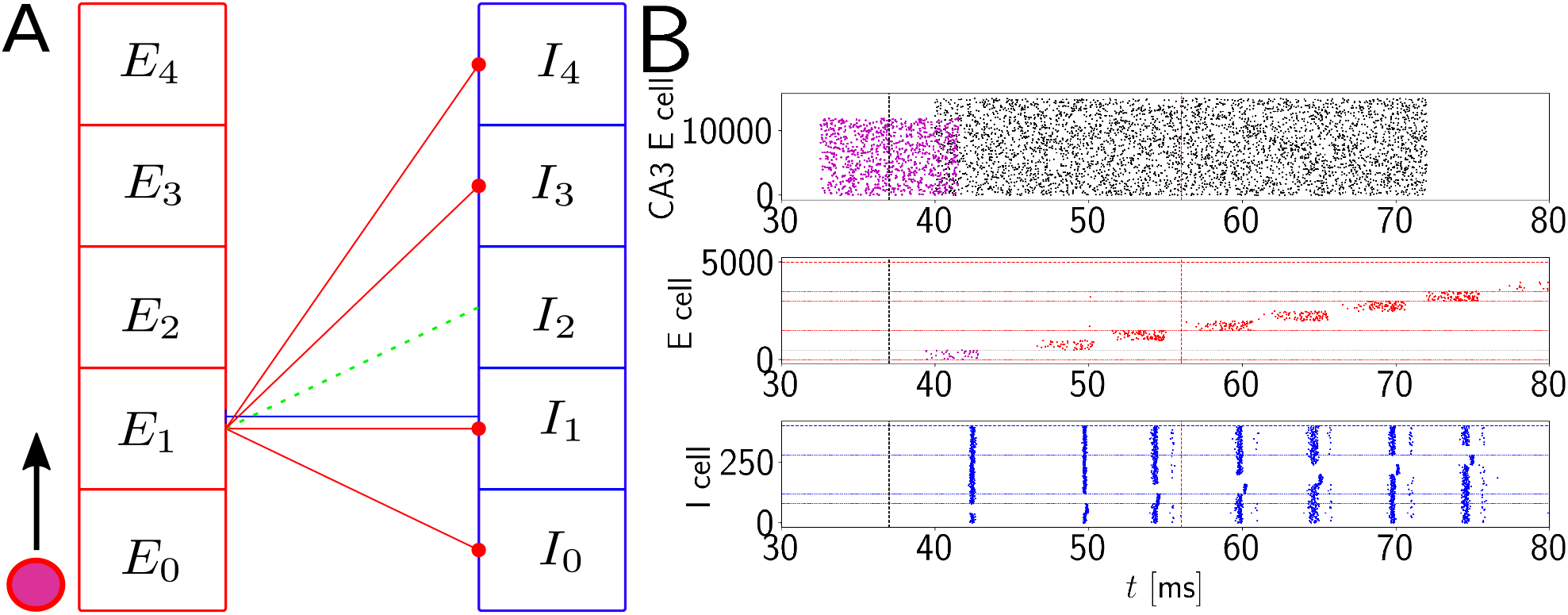
Schematic network connectivity for replay events induced by pulse and gap coding of E cells and I cells. The sequence replay is based on subsequent disinhibition of a group of E cells and the resulting excitation of all but one group of I cells, which in turn disinhibits another group of E cells. **A:** Connectivity scheme. Both the E cells and the I cells are grouped into *K* + 1 groups (here: *K* = 4 was chosen for simplicity, in the simulations below, *K* = 9). Group *E*_1_ excites all groups of I cells except *I*_2_ (green dashed line). Similarly, group *E_k_* excites all groups *I_k_*, but not *I*_*k*+1_. Group *I_k_* projects on a single group *E_k_* for all *k* = 0*, …, K* − 1, that is, group *I*_0_ inhibits group *E*_0_, group *I*_1_ inhibits group *E*_1_, and so on. All connections relevant for the sequence generation are only shown for group *E*_1_ and *I*_1_. I-to-I connections are not shown. A ripple event is triggered by the initial stimulation of group *E*_0_ and the ripple event progresses in the direction of the black arrow. The subsequent activation by CA3 lets E neurons spike when they are not inhibited. *E*_0_ activates all I neurons except *I*_1_. Since *I*_1_ inhibits all E cells except group *E*_1_, these become active due to the random, unstructured CA3 input. This excites all I cells except those forming group *I*_2_. The resulting gap in inhibition generates a pulse in E cell group *E*_2_ on the next ripple wave etc. **B:** Example realization in a network with *N_E_* = 5000 excitatory cells and *N_I_* = 400 inhibitory cells generating replay together with an oscillation of frequency *f* ≈ 190 Hz. Group *E*_0_ receives input from the purple CA3 population consisting of 12000 neurons. The remaining groups of excitatory cells receive unstructured input from *N_E,CA3_* = 15000 different excitatory CA3 spike trains shown in black. About seven E (I) groups are sequentially (de)activated, before the sequence terminates due to the termination of CA3 input. Parameters as in Fig 8, except those listed in SI section Parameters of the replay model.

Fig 10 shows that the replay by alternating excitatory pulse and inhibitory gap coding proposed for CA1 is robust, by changing the number of excitatory and inhibitory cells *N_E_* and *N_I_*. The replay occurs together with ripples in the ripple frequency range (at about 190 Hz) for intermediate values of *N_E_* and *N_I_*. To achieve such robust replay, we adjusted some of the parameters of the model compared to model 3, see SI section Parameters of the replay model for a list of all changed parameters. In particular, the peak inhibitory synaptic conductance on E cells is increased so that at the same holding potential as previously, each inhibitory spike now generates a hyperpolarization of 1.35 mv instead of 0.83 mV. This modification is biologically plausible given recent *in vivo* experiments [30], but also consistent with older studies [84]. Further, we chose the parameters of the dendritic spike strength distribution such that E cell spiking can be effectively suppressed by I cell spiking, i.e. we avoid large values for the peak dendritic current. Concretely we set the mean of the peak dendritic current distribution slightly above the value required for the generation of one spike from rest (see Fig 2), and we truncate the distribution at 4 nA as described above.

**Fig 10.**
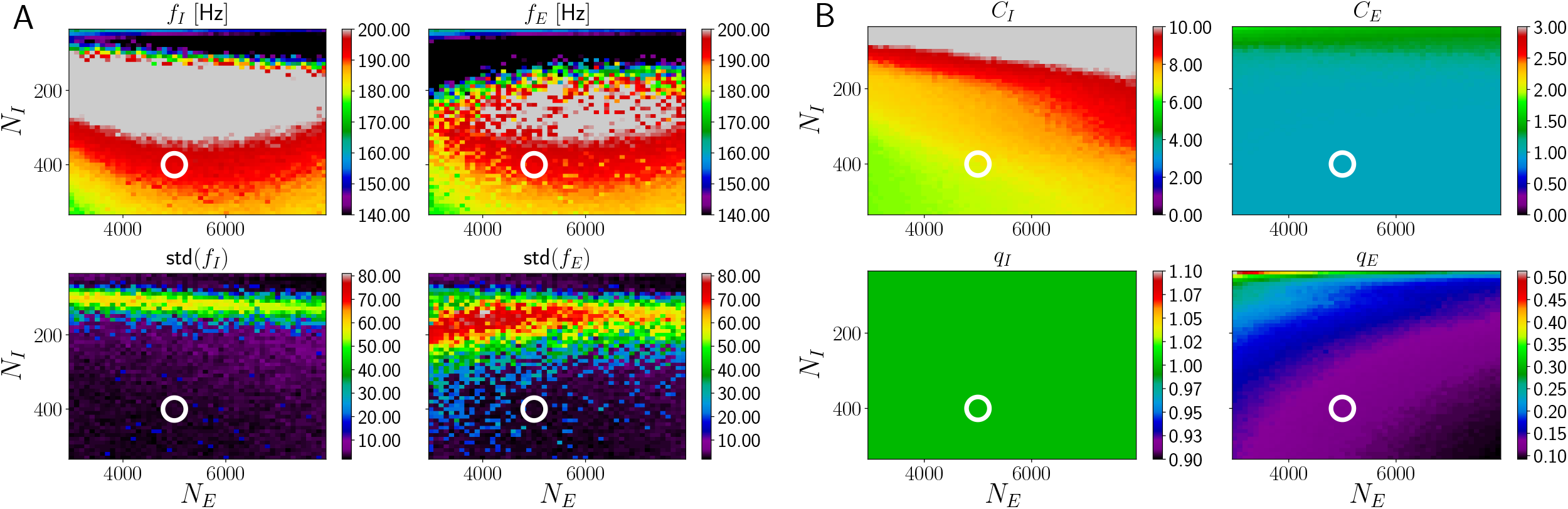
Robustness of replay events and accompanying ripple frequency oscillations. For a broad range of E and I cell population sizes, replay events accompanied by HFOs in the ripple range are robustly generated. E cells participating in such an SPW/R event mostly contribute only one spike, while all I cells participate and spike in nearly every ripple. The layout of the figure is similar to Fig 4; the changed variables are now the number of E cells *N_E_* and the number of I cells *N_I_* in a network. The populations are split into *K* + 1 = 10 groups forming chains according to alternating pulse and gap coding, as in Fig 9. A: Network frequencies and standard deviations. B: Number of spikes per active neuron and fraction of active neurons. Robust replay is observed in the upper ripple frequency range (at about 190 Hz), red in top panel of A. The white circle indicates the parameters of Fig 9 B, (*N_E_, N_I_*) = (5000, 400).

As in Fig 8, the I-to-E synaptic latency is kept at 0.5 ms. This comparably small value is helpful for robust replay accompanied by HFOs in the ripple range: a simulation with increased latency is shown in the SI, Fig 29; there, the HFOs are less robust (increase of the standard deviation across realizations) and the maximal frequency decreases to approximately 170 Hz for the E population. We conclude that fast and strong inhibition is beneficial for our proposed replay scheme.

## Discussion

In the current article, we have studied two-population models for the generation of ripples and sequences in the hippocampal region CA1. In these models E and I neurons interact to generate the ripple rhythm possibly together with sequential activity. This is motivated by recent *in vivo* experiments, which have shown that both the local PC and the local PV+BC populations in CA1 contribute to generating ripple oscillations [22]. Further motivation comes from the observation that the connectivity from local PCs to PC+BCs and back is high. Our models are constrained by biologically plausible connectivity and by the fact that during ripple oscillations, basically all PV+BCs spike at high frequency, i.e. on nearly every ripple wave, while the PCs spike sparsely, contributing typically only one or two spikes.

We observe that two-population models receiving temporally broad CA3 sharp wave input either generate ripple frequency oscillations or sparse spiking of E neurons, but not both. Sparse spiking of E neurons can be reached if there is a strong CA3 projection to the I neurons, such that the oscillations are effectively generated by the I population alone, which seems inconsistent with experimental findings [22]. We have thus explored in more depth another idea, namely that the sparse spiking of E cells in CA1 originates from temporally sparse, short and strong inputs from CA3. Different CA1 PCs receive these inputs at different times, their density is highest near the peak of the sharp wave. We propose that such sparse inputs might originate from dendritic spikes in the apical dendrites, which are elicited when enough spikes arrive from CA3 within a short time window [65, 67].

The inclusion of pyramidal cells and the condition of their sparse firing renders the generation of high frequency oscillations challenging compared to models where the oscillation is basically generated by I neurons only. This is on the one hand because the additional E-to-I and I-to-E loop generally slows down the network oscillation frequency [37]. On the other hand, we find that simply increasing the level of external excitation to increase the oscillation frequency usually results in too much E cell spiking, which renders the models unsuitable to describe ripple oscillations. Previous two-population models for HFOs have assumed connection probabilities[27, 29, 37, 117] and/or spiking dynamics[25, 29, 37, 117] that do not fit the situation in CA1 [1, 20, 57]. Characteristic for the spiking activity during CA1 SPW/Rs is that they are a mixture of two often considered oscillation types: strongly synchronized [36] and weakly synchronized oscillations [25, 27, 37]. The E cell population is weakly synchronized; the average single-cell firing frequency during a ripple is much lower than the population frequency. The I cell population (consisting of PV+BCs) is strongly synchronized, with every I cell spiking at nearly each individual ripple wave. In the current article, we aimed at taking into account both the experimental knowledge on CA1 connectivity and the characteristics of the E and I cell spiking activity during SPW/Rs.

How realistic are our assumptions concerning the generation of dendritic spikes? Apical dendritic spikes have been directly observed during SPW/Rs [86]. It is difficult to estimate the number of synaptic inputs required to generate dendritic spikes, estimates range “anywhere from a handful to dozens of inputs” [118]. Detailed multicompartmental models of morphologically reconstructed neurons suggested that at least ~50 synaptic inputs arriving within 1 – 3 ms on a small part of the apical dendrite of CA1 neurons are required to elicit dendritic sodium spikes [65, 67]. In our model 3, we assume that already 5 spikes arriving within 2 ms are sufficient for dendritic spike generation. This is similar to the number of inputs required for spike generation in basal dendrites of CA1 pyramidal cells [43, 44, 66]. Increasing the threshold for dendritic spike generation while keeping the number of afferent inputs and the dendritic integration window *w_D_* constant necessitates an increase of the afferent rate. In our model 3, increasing the dendritic spike threshold from 5 to 50 would require an increase of *r*_0_ in Eq 7 from 8 Hz to more than 100 Hz to approximately maintain the same number of dendritic spikes. This value is clearly too high as a discharge rate for a typical CA3 cell during sharp waves [64] (but values larger than 10 Hz are possible [60]). The inputs from CA3 to CA1 may, however, be clustered such that sufficient coincident inputs impinge on an apical dendrite. Our result might also indicate that a smaller number of coincident inputs than previously estimated is required to elicit dendritic spikes. Further, dendritic spikes relevant during SPW/Rs may be less sensitive to synchrony than assumed in our model, i.e. their effective dendritic integration window may be longer. Such dendritic spikes might then be NMDA or calcium spikes [71, 86, 119–121]. CA3 cell bursting [60] might in principle also contribute to dendritic spike generation, for example because asynchronous but overlapping bursts generate synchronous spike inputs. Experimentally found ISIs within individual CA3 bursts [60] are, however, typically larger than the sodium dendritic spike integration window [65, 66]. Finally, basal dendrites could generate dendritic spikes as incorporated in our model, since they receive inputs from CA3 PCs besides that from CA1 PCs [15]. Particularly promising candidates are the recently discovered axon carrying dendrites, from which the axon emanates in many CA1 PCs [69]. These dendrites are particularly excitable, generate strong dendritic spikes and have a high impact on action potential generation. Ref. [22] found that stimulation of the local E CA1 population can excite ripple oscillations. Such stimulation may generate dendritic spikes in the basal dendrites of postsynaptic CA1 E cells [43, 44], which could lead to ripples in a similar manner as in our model 3 (or in the manner described in ref. [43, 44]).

PC spiking during SPW/Rs in our model 3 is generally very sparse, the majority of PCs contributes one to two spikes, as observed experimentally. In the model without sequence replay a few CA1 PCs spike more than 3 times during a SPW/R event (Fig 7). Such bursting is consistent with experiments[39, 60, 100, 101]. The bursts are generated in our model because the lognormal distribution of the peak dendritic currents across neurons (Eq 2) contains larger values with non-negligible probability. Additional simulations with truncated lognormal or Gaussian distributions with less bursting and simulations with a long somatic refractory period, which completely prevents multiple spiking, can generate ripple range HFOs. This shows that bursts are not necessary for the generation of HFOs in the ripple frequency range in our model.

HFOs in the ripple range occur in model 3 (Fig 8), at or before the border of stability where standard deviations across realizations increase. This happens because a certain critical amount of externally supplied excitation (quantified by the parameters of the peak dendritic current in model 3) is needed to generate HFOs in the ripple range. Below this level, gamma and high gamma oscillatory states are reached. Beyond this level, E cell spiking cannot be organized into distinct ripple waves by PV+BC spiking anymore; the E cells are permanently active and show little oscillatory modulation in their spiking.

A prominent property of the hippocampus is that it generates sequences of activity. These may replay previously imprinted sequential experience [101, 103, 104], serve as a backbone to store episodic memories [122, 123] or generally provide a sequential reference frame for sensory experience and brain activity [124]. Sequence generation is often assigned to CA3 because of its prominent, albeit sparse, recurrent excitatory connectivity [122, 123]. However, CA1 also generates sequences on its own [39]. Earlier work proposed that the highly sparse recurrent excitatory connectivity in CA1 could underlie sequence generation, since it may be highly structured and amplified by basal dendritic spikes [43, 44]. Here we propose a different class of models for sequence generation in CA1: two-population models. In such models the sequences are generated cooperatively by the E and I population. We conceptually propose two such models, one where the sequence generation depends on the prominent E-to-I neuron and I-to-E neuron connectivity and another one where also the similarly prominent I-to-I neuron connectivity is important. We explicitly implemented the first one of these concepts, since it is more plausible for CA1. It is based on alternating pulse and gap coding of the E and I neuron populations, such that both the E and the I neuron firing patterns together generate the sequence. This is different from the classical view that sequence generation in neural networks depends mainly on the excitatory connectivity between E neuron groups like in synfire chains [125–128], while inhibition prevents pathological activity, allows gating and introduces competition between sequences [129–135]. Specifically, inhibition of the embedding network and of previously active groups by propagating synfire chain activity prevents pathological, strong increases of overall network spiking activity [129, 133]; the stability of propagation along the synfire chain can then be improved by additional inhibitory neuron groups: their sparse feedforward activation leads to disinhibitory removal of excessive inhibition from specific excitatory groups [135]. We have shown that two-population based sequential replay is compatible with filtering of CA3 input by dendritic spikes, as proposed in model 3, and that it generates high frequency oscillations with sparse E cell firing. We expect that the latter holds also for different forms of input from CA3 such as those explored in model 1.

Two-population based sequential structures may be present in further areas beyond the hippocampal area CA1. They may consist of discrete groups or be continuous and they may be preexisting or learned during experience. It is an interesting direction of future research to determine how their spontaneous formation or learning may take place through the interplay of excitatory and inhibitory synaptic plasticity [44, 136–138].

To conclude, based on neurobiological knowledge on the hippocampal regions CA1 and CA3, their single neurons and their dynamics, we have shown how CA3 drive and two-population interactions in the region CA1 may lead to the SPW/R population pattern, with its characteristic high frequency oscillations and sparse E and dense I cell firing. For the associated sequential replay we have developed a model that is consistently based on two-population interactions, specifically on E pulse and I gap coding.

## Supporting information

### Parameters

**Table 1.** Parameters of E cells.

|  |  |  |  |
| --- | --- | --- | --- |
| $C_E$ | 275 pF | $g_l^E$ | 25 nS |
| $E_{\text{rest}}^E$ | -67 mV | $E_{\text{exc}}^E$ | 0 mV |
| $E_{\text{inh}}^E$ | -68 mV | $(N_E)$ | (12000) |
| $V_{\text{thresh}}^E$ | -50 mV | $V_{\text{reset}}^E$ | -67 mV |
| $\tau_{\text{ref}}^E$ | 2 ms | | |

**Table 2.**
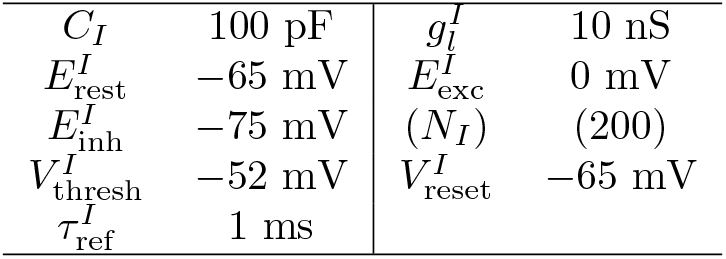
Parameters of I cells.

|  |  |  |  |
| --- | --- | --- | --- |
| $C_I$ | 100 pF | $g_l^I$ | 10 nS |
| $E_{\text{rest}}^I$ | -65 mV | $E_{\text{exc}}^I$ | 0 mV |
| $E_{\text{inh}}^I$ | -75 mV | $(N_I)$ | (200) |
| $V_{\text{thresh}}^I$ | -52 mV | $V_{\text{reset}}^I$ | -65 mV |
| $\tau_{\text{ref}}^I$ | 1 ms | | |

**Table 3.** Parameters of synapses on E cells.

|  |  |  |  |
| --- | --- | --- | --- |
| $g_{\text{exc,peak}}^E$ | 0.9 nS | $g_{\text{inh,peak}}^E$ | 9.0 nS |
| $\tau_{\text{exc},d}^E$ | 1.8 ms | $\tau_{\text{exc},r}^E$ | 0.5 ms |
| $\tau_{\text{inh},d}^E$ | 2.0 ms | $\tau_{\text{inh},r}^E$ | 0.4 ms |

**Table 4.** Parameters of synapses on I cells.

|  |  |  |  |
| --- | --- | --- | --- |
| $g_{\text{exc,peak}}^I$ | 3.0 nS | $g_{\text{inh,peak}}^I$ | 5.0 nS |
| $\tau_{\text{exc},d}^I$ | 1.2 ms | $\tau_{\text{exc},r}^I$ | 0.5 ms |
| $\tau_{\text{inh},d}^I$ | 1.2 ms | $\tau_{\text{inh},r}^I$ | 0.45 ms |

The latencies *τ_l_* are fixed at 1 ms for all four types of synapses if not mentioned otherwise. At the beginning of a simulation, the neurons are initialized with voltages drawn from Gaussian distributions centered around their respective resting potentials with standard deviation 0.1 mV.

### Synaptic dynamics in CA1

Fig 11 displays excitatory and inhibitory postsynaptic potentials of the four synapse classes present in CA1 with the standard parameters used in our models.

**Fig 11.**
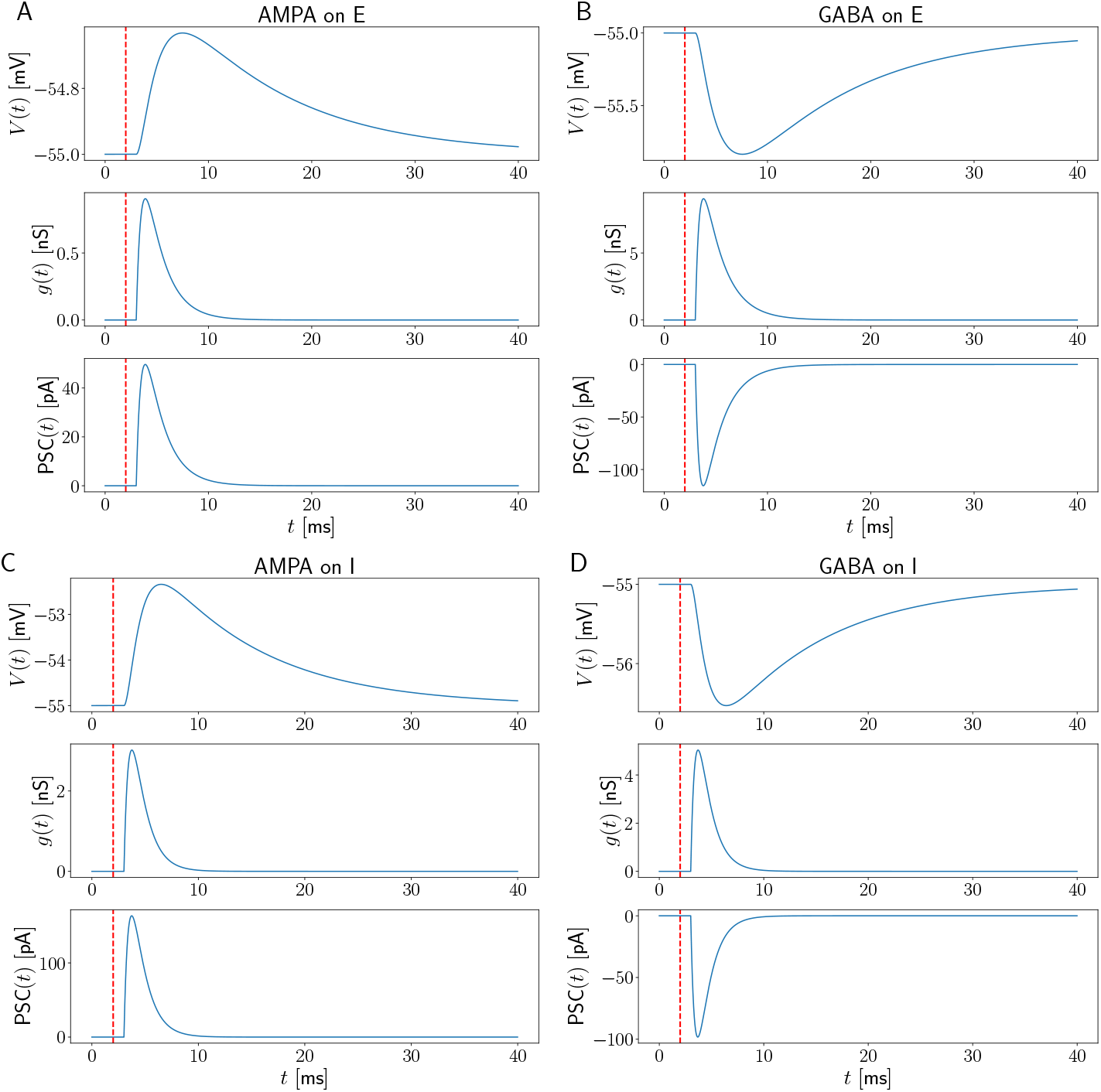
Synaptic dynamics in CA1. At *t* = 2 ms (red vertical dashed line) a presynaptic spike occurs which triggers a conductance change *τ_l_* = 1 ms later. Membrane voltage (top panel), corresponding synaptic conductance (middle panel) and post-synaptic current (PSC, bottom panel) for A: E-to-E synapses, B: I-to-E synapses, C: E-to-I synapses, D: I-to-I synapses. Holding potential: −55 mV. *σ_n_* = 0 mV. Values for time constants and peak conductances can be found in the SI, section Parameters.

### Control simulations for model 1

In Figs 12 - 14, the displayed frequency range for *f_I_* and *f_E_* is enlarged to [100, 200] Hz to yield a broader overview. The plot layout is as in Fig 4.

#### Higher I-to-E, I-to-I connectivity and broader sharp waves in model 1

**Fig 12.**
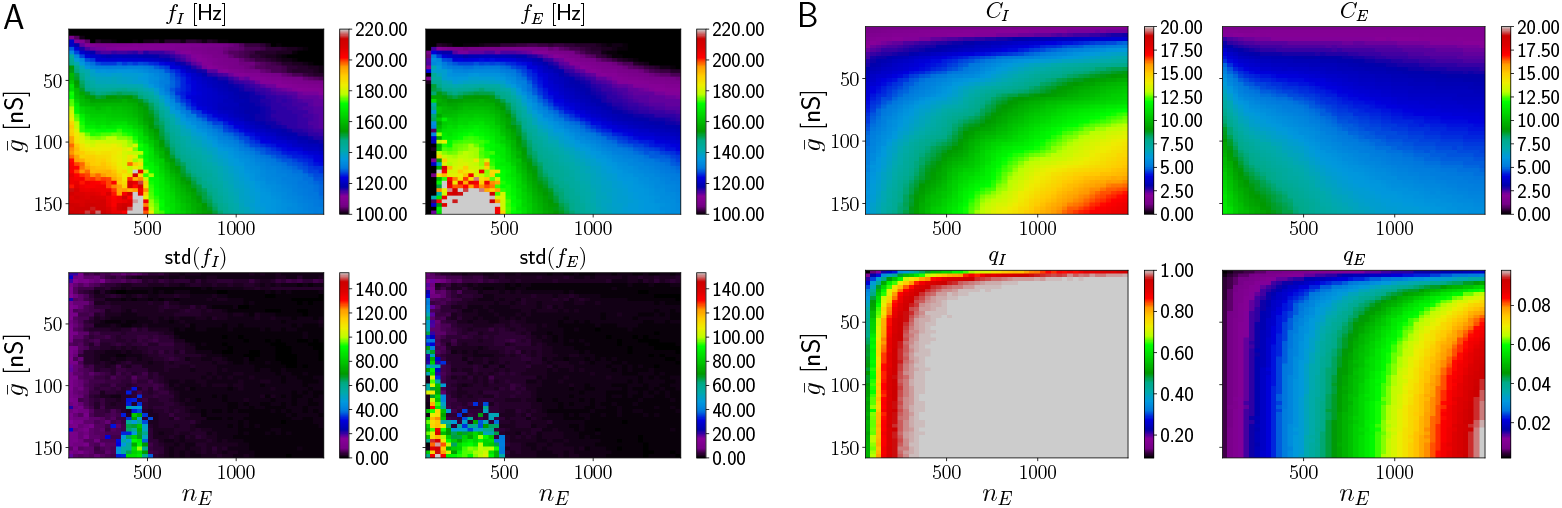
HFOs in networks with temporally homogeneous excitation of E cells and higher I-to-E connectivity. Parameters as in Fig 4, except higher *p_IE_* = 0.2.

**Fig 13.**
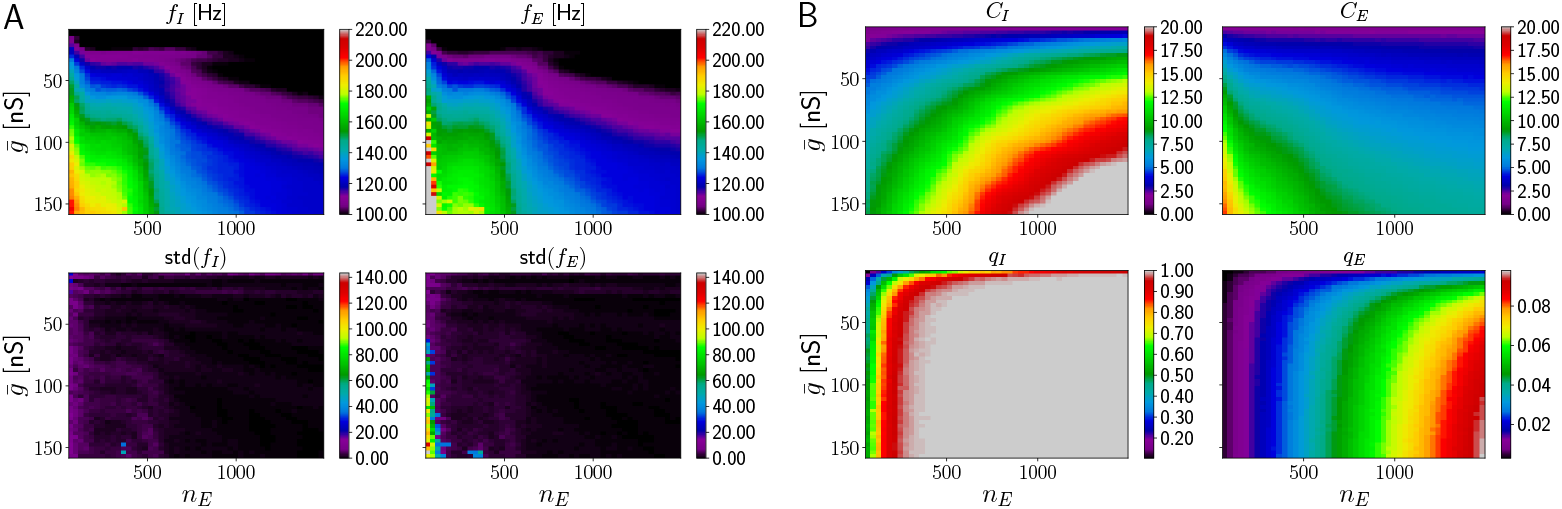
HFOs in networks with temporally homogeneous excitation of E cells and higher I-to-E connectivity as well as broader sharp waves. Parameters as in Fig 4, except higher *p_IE_* = 0.3 and larger width of sharp waves *σ_g_* = 15 ms.

**Fig 14.**
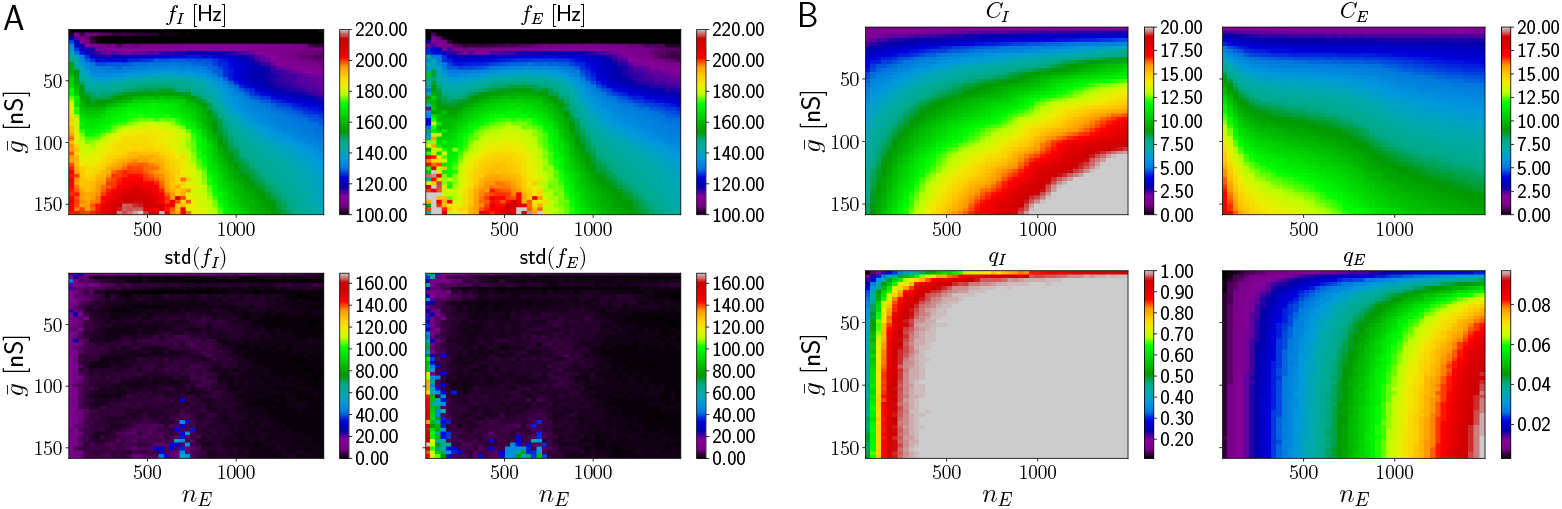
HFOs in networks with temporally homogeneous excitation of E cells and higher I-to-I connectivity, higher I-to-E connectivity and broader sharp waves. Parameters as in Fig 4, except higher *p_II_* = 0.3, *p_IE_* = 0.2 and larger width of sharp waves *σ_g_* = 15 ms.

#### Strong feedforward excitation to I cells in model 1

**Fig 15.**
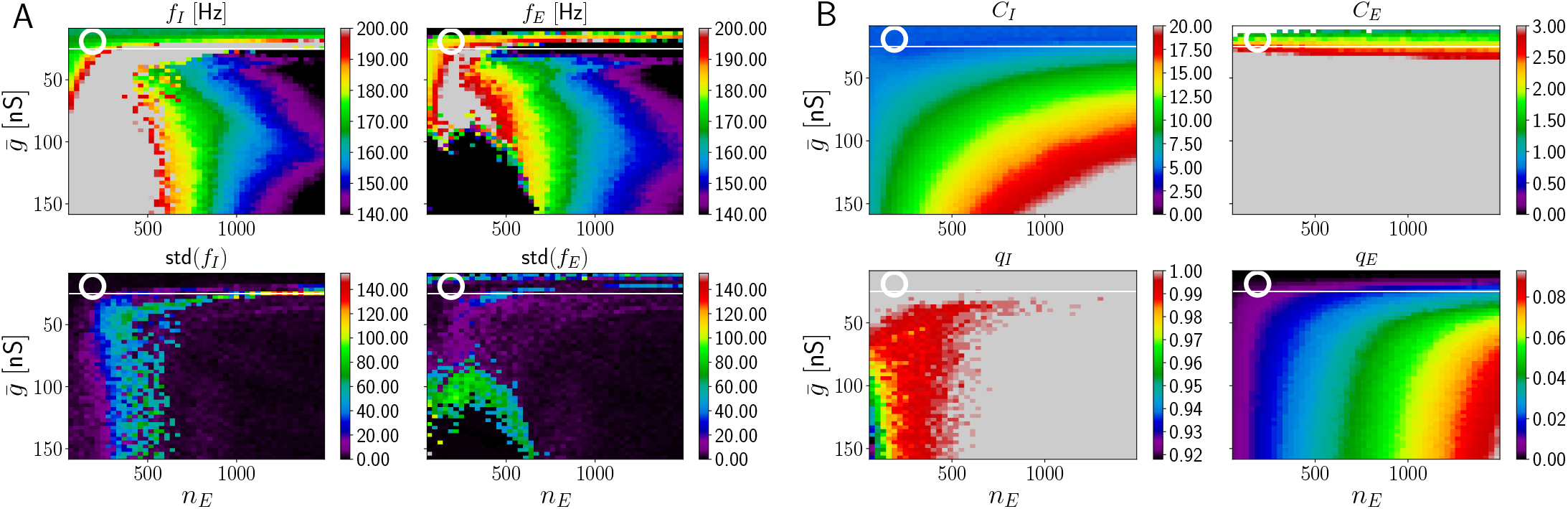
HFOs in networks with temporally homogeneous excitation of E cells and strong feedforward excitatory drive to I cells. A, upper subpanel: Frequency of I and E population activity oscillations *f_I_*, *f_E_* as a function of mean drive 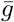 and number of excited E cells *n_E_* (average taken across different network realizations). Lower: Corresponding standard deviations across network realizations, std(*f_I_*) and std(*f_E_*). B, upper subpanels: mean number of spikes per active inhibitory (excitatory) neuron *C_I_* (*C_E_*). Lower: fraction of active neurons (*q_I_* and *q_E_*). The range for *f_I_* and *f_E_* is [140, 200] Hz. The ranges for *C_I_* and *C_E_* are [0, 20] and [0, 3], respectively. Frequency values below (above) this range are indicated in black (gray). The white horizontal line is located at 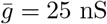 roughly delimiting the region of sparse E firing. The white circle is at 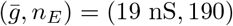 (parameters of Fig 16). Parameters as in Fig 4, except for drive of I cells: Each PV+BC is driven by a conductance as in Eq 6, with width *σ_g_* = 10 ms and fixed amplitude 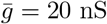

**Fig 16.**
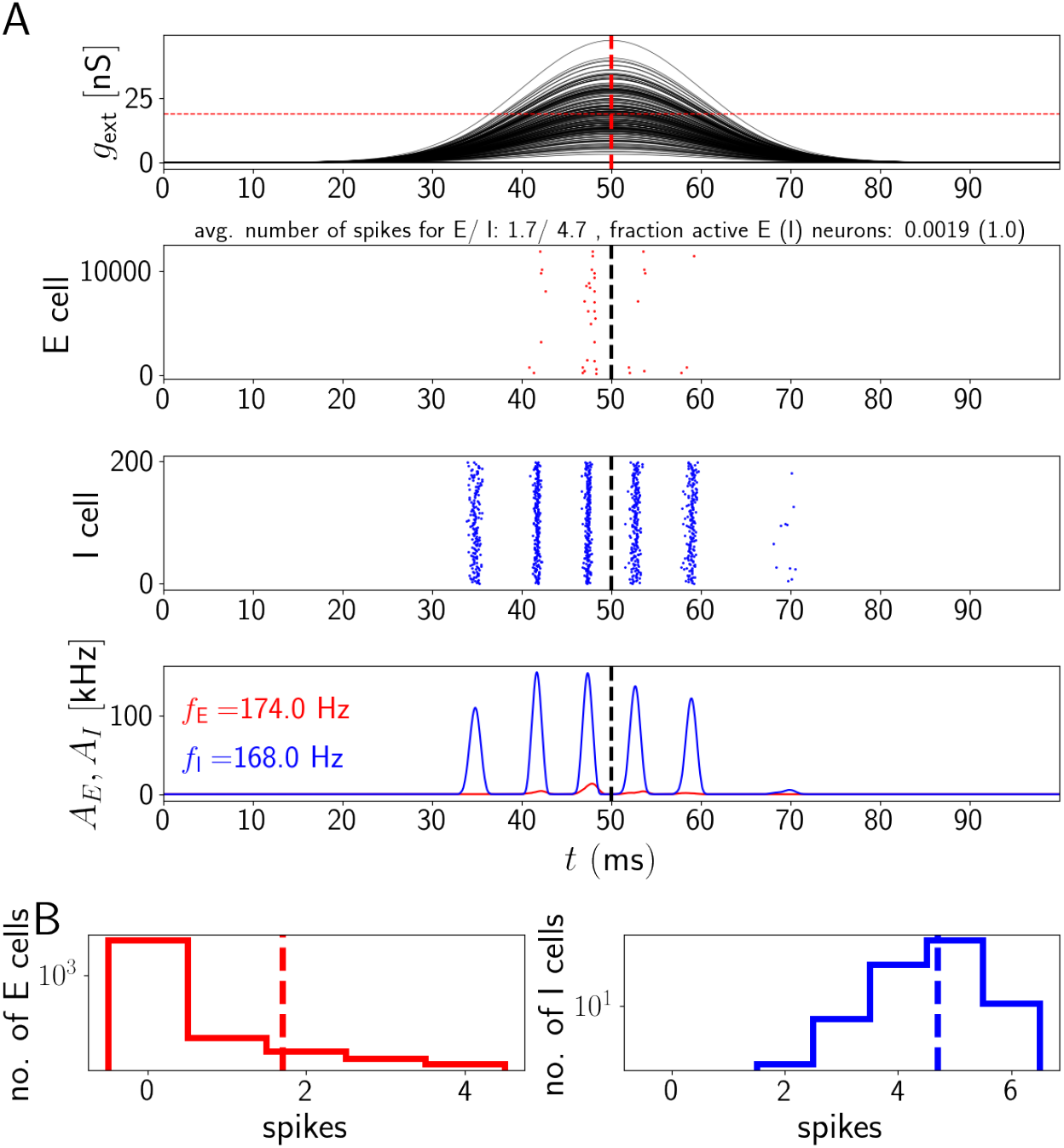
Network activity for temporally homogeneous excitation of E cells and strong feedforward excitatory drive to I cells. A: Sharp wave time courses for E cells, spike rastergrams and network activities. The I cells receive sharp wave input with fixed amplitude 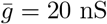 and *σ_g_* = 10 ms. The horizontal red dashed line is located at the mean 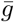 of the sharp wave amplitudes. B: Histograms of spike counts for E (left) and I (right) cells on a logarithmic scale. Parameter values as in Fig 15, parameters of E cell drive are 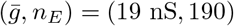.

### Control simulations for model 2

In Figs 17 - 20, the frequency range for *f_I_* and *f_E_* is set to [100, 200] Hz. The plot layout is as in Fig 6.

#### Broader pulses in model 2

**Fig 17.**
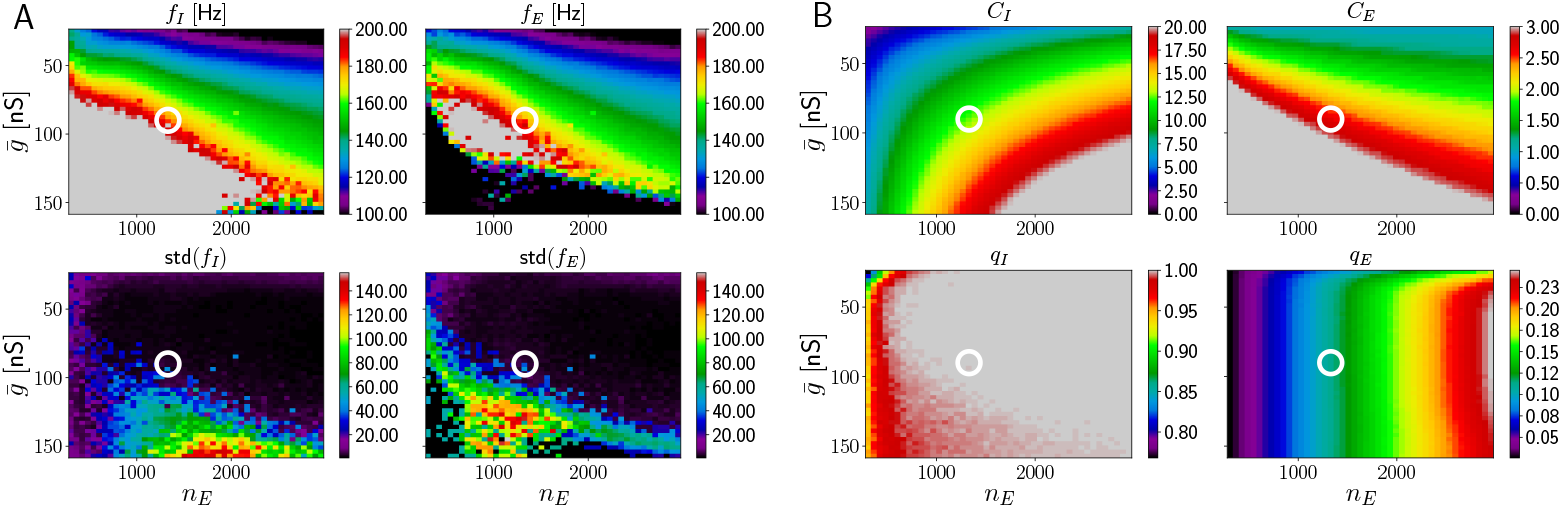
HFOs in networks with temporally inhomogeneous excitation of E cells and broader input pulses. Sparse spiking of E cells is largely lost in the parameter region where ripple frequency HFOs occur. The white circle is located at 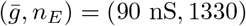. In this region, HFOs exist, but E cells spike more than two times on average, which is not realistic. Parameters as in Fig 6, except *σ_g_* = 5 ms.

#### Larger I-to-E, I-to-I connectivity and broader spread of individual input pulses in model 2

**Fig 18.**
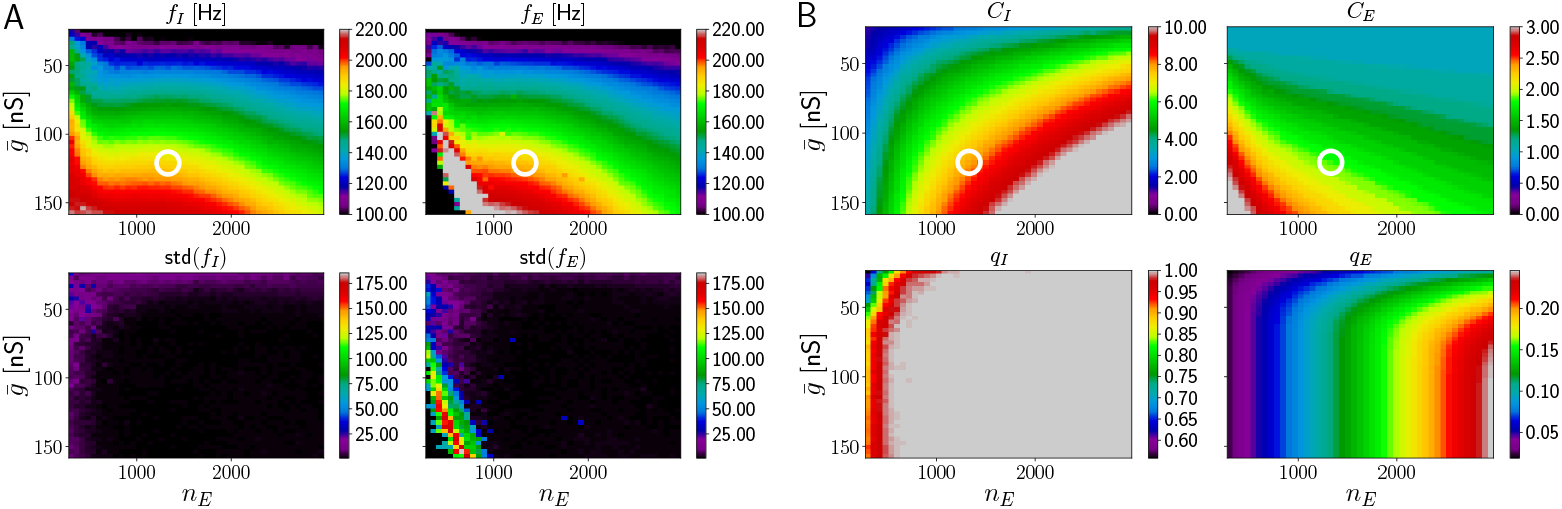
HFOs in networks with temporally inhomogeneous excitation of E cells and larger I-to-E and I-to-I connectivity. The white circle is located at 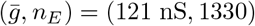. It indicates a region where ripple frequency oscillations and sparse E cell spiking still occur. Parameters as in Fig 6, except *p_II_* = 0.3 and *p_IE_* = 0.2.

**Fig 19.**
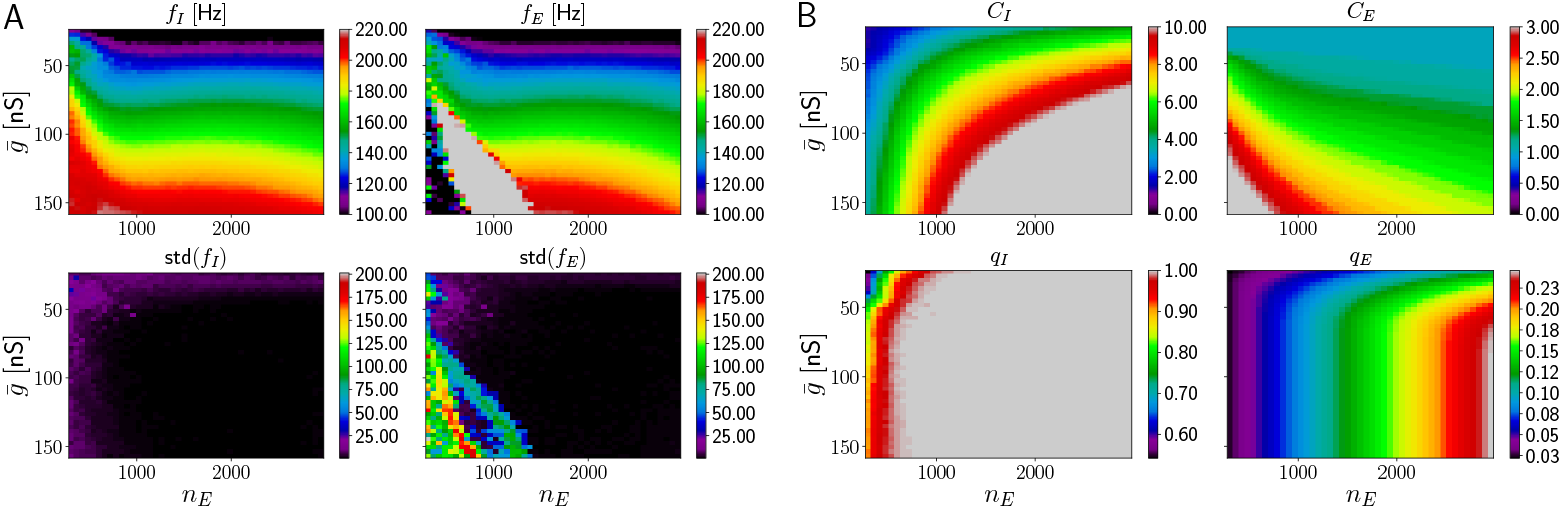
HFOs in networks with temporally inhomogeneous excitation of E cells and broader spread of individual input pulses as well as larger I-to-E and I-to-I connectivity. Parameters as in Fig 6, except *σ_t_* = 15 ms as well as *p_II_* = 0.3 and *p_IE_* = 0.2.

**Fig 20.**
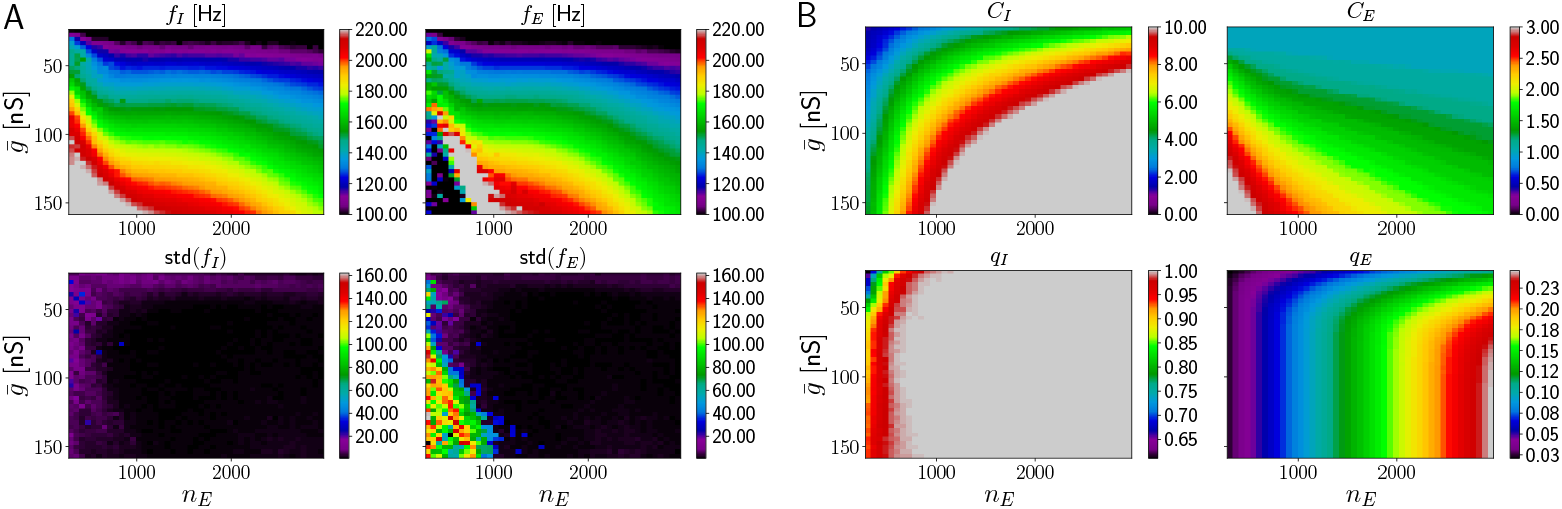
HFOs in networks with temporally inhomogeneous excitation of E cells and broader spread of individual sharp wave peaks as well as higher I-to-E connectivity. Parameters as in Fig 6, except *σ_t_* = 15 ms and *p_IE_* = 0.2.

### Control simulations for model 3

In Figs 21 - 25, the frequency range for *f_I_* and *f_E_* is set to [100, 200] Hz. The plot layout is as in Fig 8. The white circle indicates regions where HFOs in the ripple range are generated and E cells fire sparsely.

#### Higher*p_IE_*, truncation of distribution for 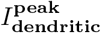, higher I-to-E synaptic latency and lower E-to-I peak conductance in model 3

**Fig 21.**
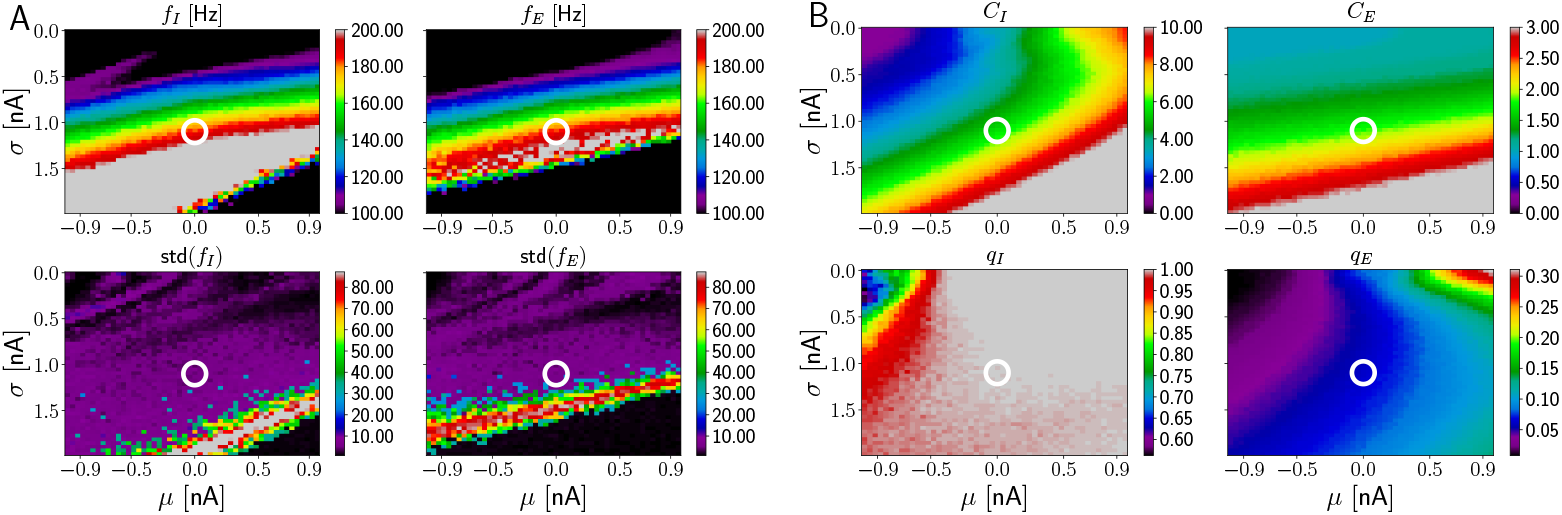
HFOs in networks incorporating dendritic excitation and larger I-to-E connection probability. The white circle is located at (*σ, μ*) = (1.1, 0.0) nA. It marks a point in the (*σ, μ*) parameter space where HFOs in the ripple range (~ 190 Hz) are generated and E cells fire sparsely. Parameters as in Fig 8, except *p_IE_* = 0.3 instead of 0.1.

**Fig 22.**
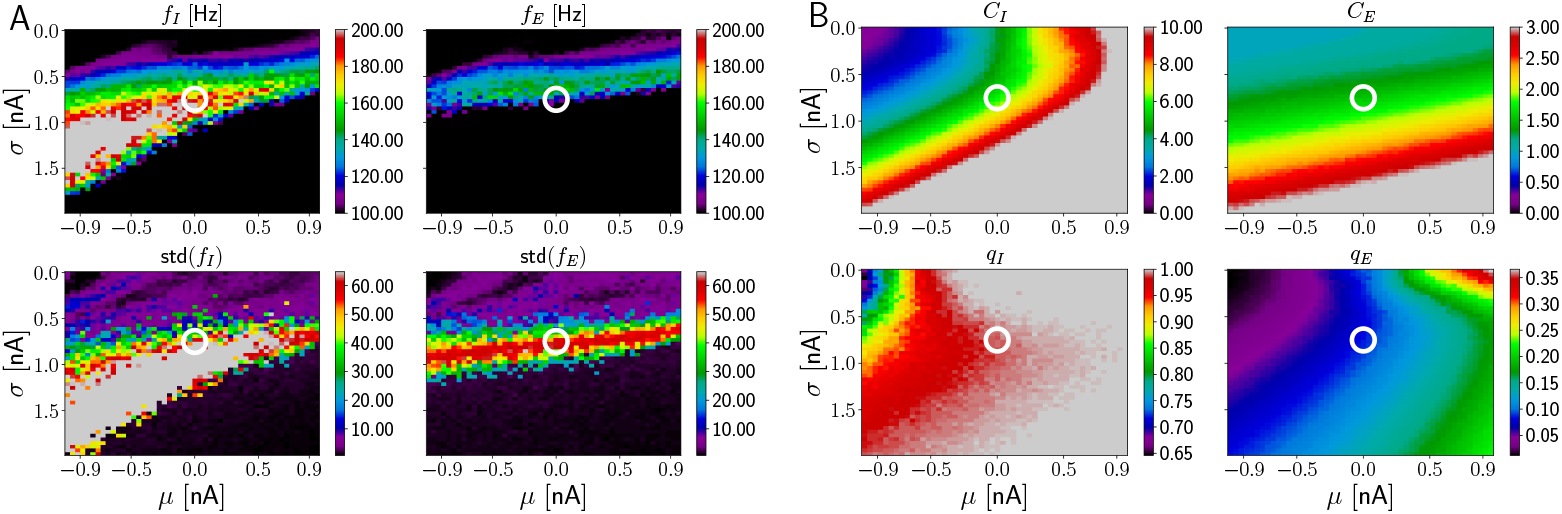
HFOs in networks incorporating dendritic excitation and higher I-to-E synaptic latency. The white circle is located at (*σ, μ*) = (0.75, 0.0) nA. Parameters as in Fig 8, except for a higher I-to-E latency *τ_l_* = 0.9 ms instead of 0.5 ms.

**Fig 23.**
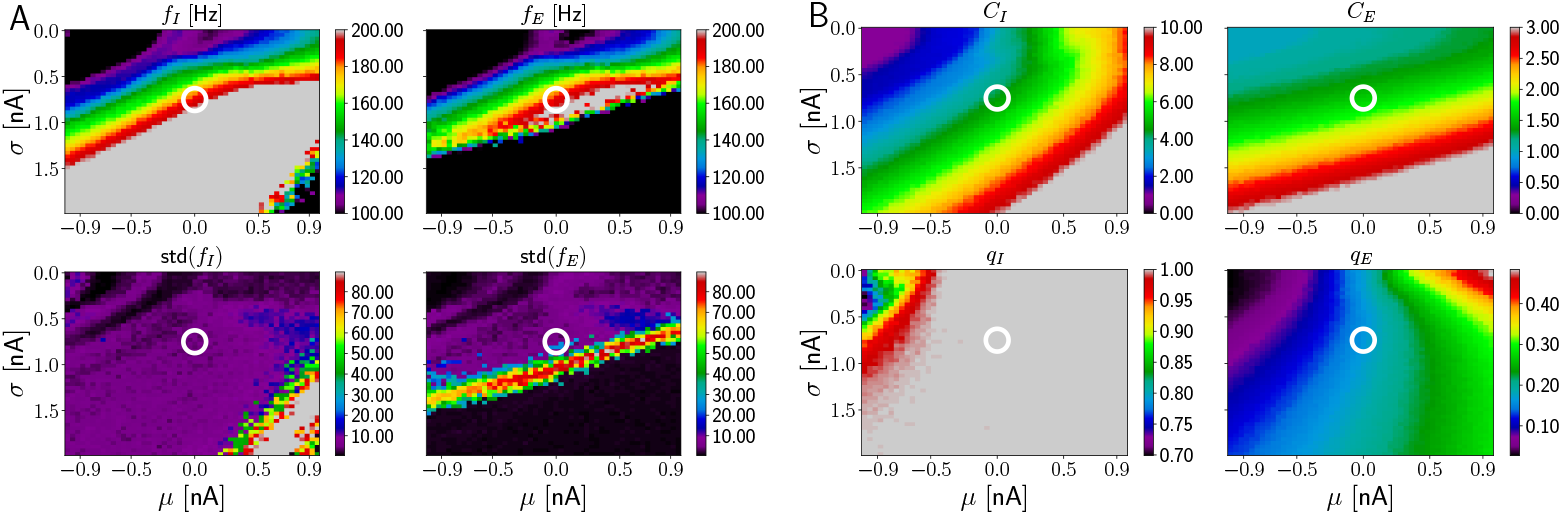
HFOs in networks incorporating dendritic excitation and lower E-to-I peak conductance. The white circle is located at (*σ, μ*) = (0.75, 0.0) nA. Parameters as in Fig 8, except for a lower E-to-I peak conductance 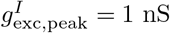 instead of 3 nS.

**Fig 24.**
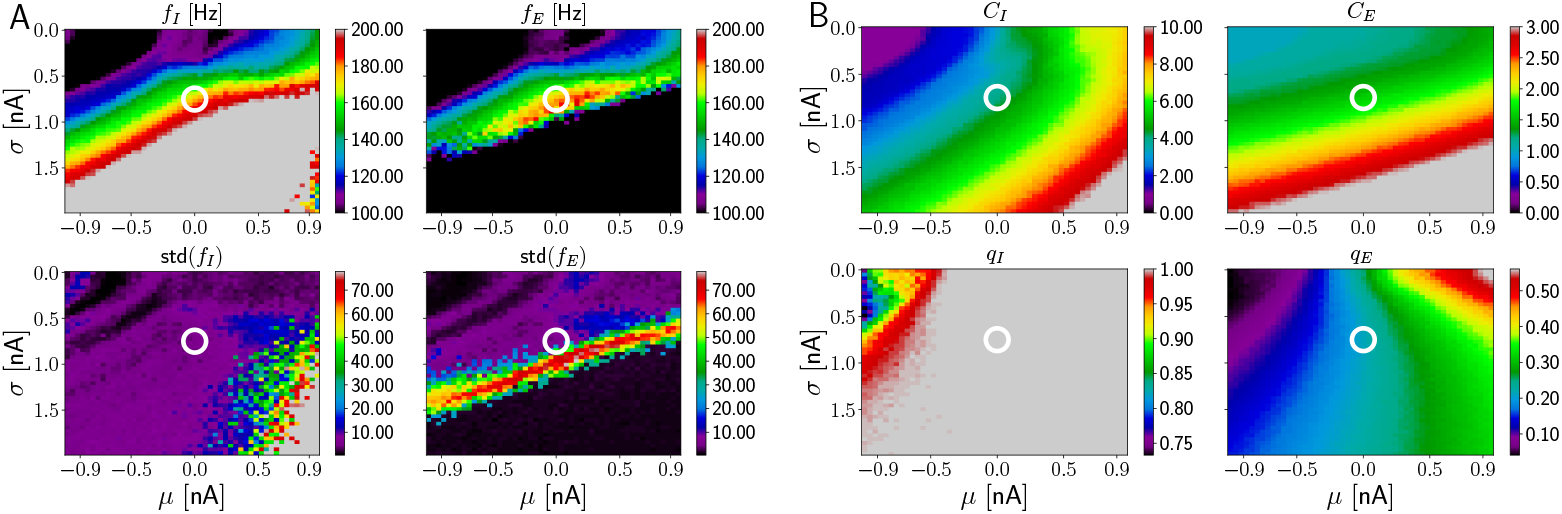
HFOs in networks incorporating dendritic excitation and higher I-to-E latency, faster E-to-I rise time and lower E-to-I peak conductance. The white circle is located at (*σ, μ*) = (0.75, 0.0) nA. Parameters as in Fig 8, except for a higher I-to-E latency *τ_l_* = 1.0 ms instead of 0.5 ms, faster E-to-I rise time 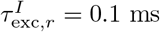 instead of 0.5 ms and lower E-to-I peak conductance 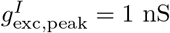 instead of 3 nS.

**Fig 25.**
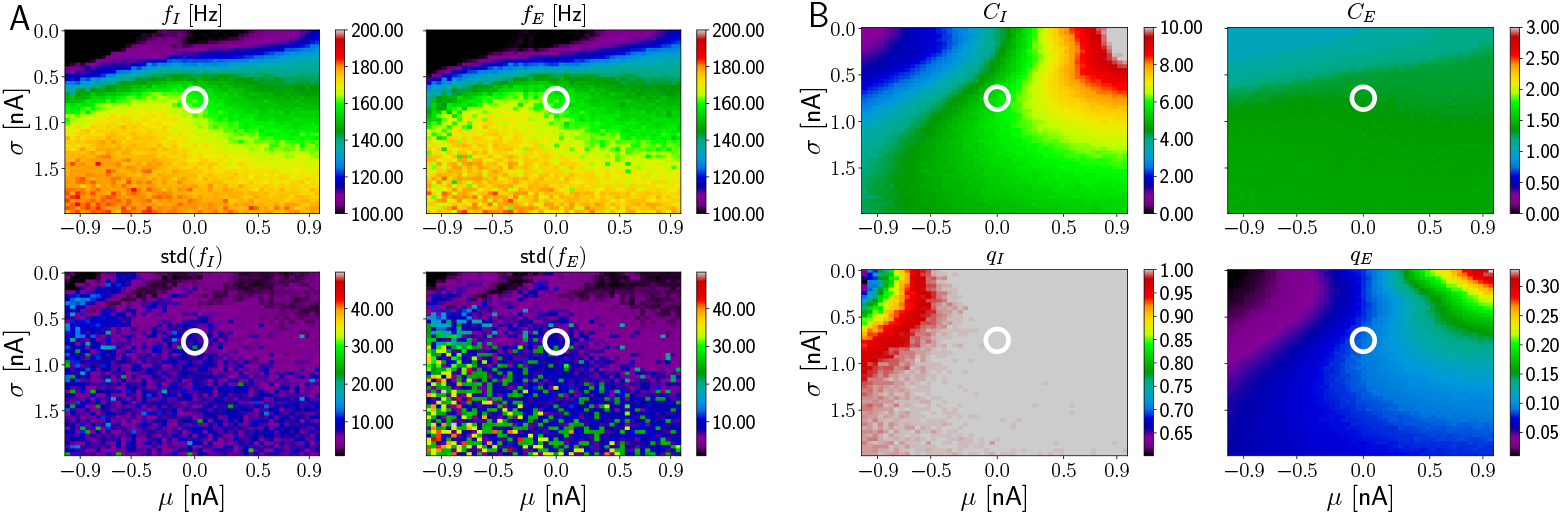
HFOs in networks incorporating dendritic excitation and a truncated distribution for the peak dendritic current. The white circle is located at (*σ, μ*) = (0.75, 0.0) nA. Parameters as in Fig 8, except for a truncation of the lognormal distribution Eq 2 at 4 nA.

**Fig 26.**
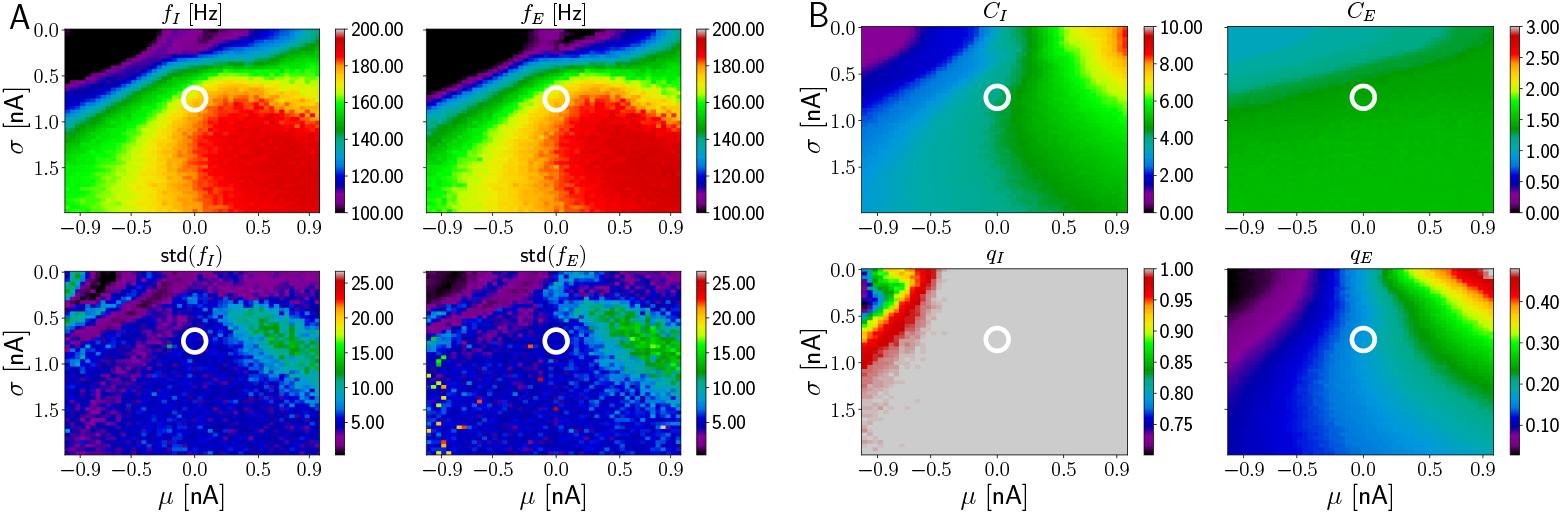
HFOs in networks incorporating dendritic excitation and a truncated re-sampled distribution for the peak dendritic current as well as lower E-to-I peak conductance. The white circle is located at (*σ, μ*) = (0.75, 0.0) nA. Parameters as in Fig 8, except for a truncation of the lognormal distribution Eq 2 at 4 nA with re-sampling of values larger than the truncation value and lower E-to-I peak conductance 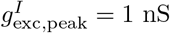 instead of 3 nS.

**Fig 27.**
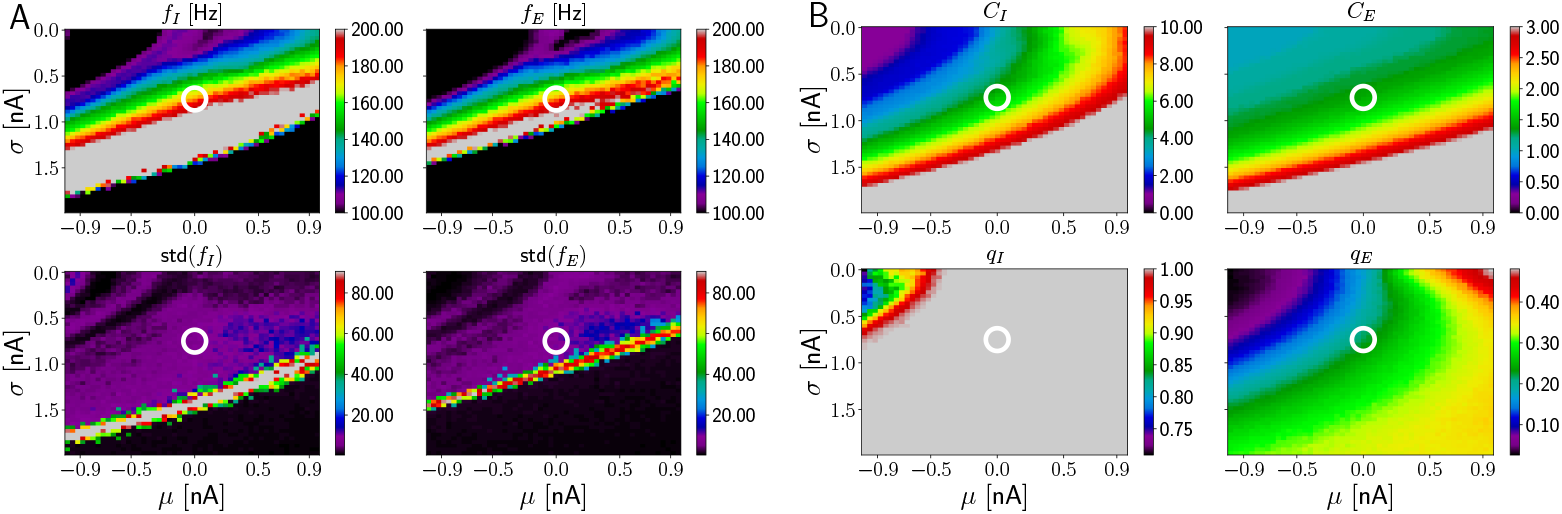
HFOs in networks incorporating dendritic excitation and a Gaussian distribution for the peak dendritic current as well as lower E-to-I peak conductance. The white circle is located at (*σ, μ*) = (0.75, 0.0) nA. Parameters as in Fig 8, except for a Gaussian distribution of the peak dendritic current 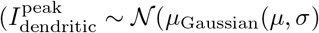, *σ*_Gaussian_(*μ, σ*)) and lower E-to-I peak conductance 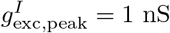 instead of 3 nS. The mean and standard deviation of theanGaussi distribution are matched to the mean and standard deviation of a lognormal distribution with parameters *μ* and 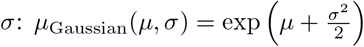 *σ*_Gaussian_(*μ, σ*) = [exp (*σ*^2^) − 1] exp (2*μ* + *σ*^2^); to allow a direct comparison with the previous figures, for parameters *μ* and *σ* are used for the plot axes.

**Fig 28.**
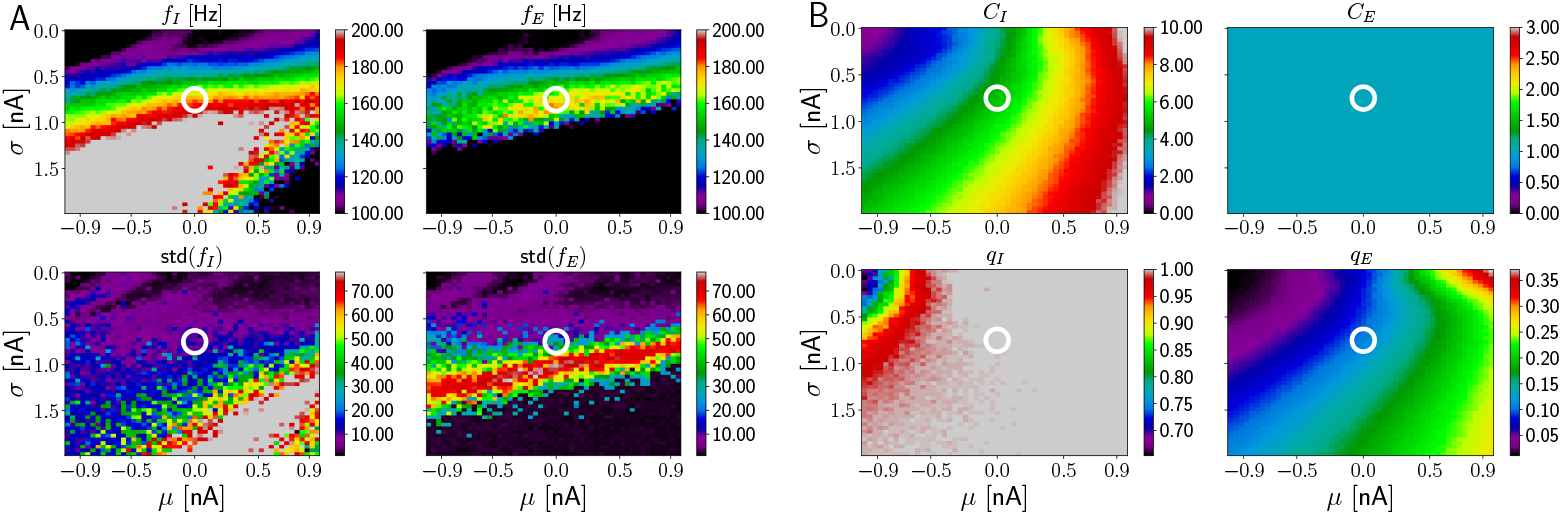
HFOs in networks incorporating dendritic excitation and a long somatic absolute refractory period for the E cells. The white circle is located at (*σ, μ*) = (0.75, 0.0) nA. Parameters as in Fig 8, except for an absolute somatic refractory period 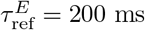. Every active E cell spikes once.

### Additional information on the replay model

#### Parameters of the replay model

The E-to-E connectivity is such that each E cell receives approximately 200 connections from other E cells. The I-to-I connectivity is such that each I cell receives 40 connections from other I cells. The CA3-CA1 excitation is exclusively via dendritic spikes, i.e. there are no small depolarizations elicited by each CA3 spike as in Fig 8. The remaining parameters are as in Fig 8, except those mentioned below. The reference values from Fig 8 are given in parenthesis.

AMPA peak conductance on I cells: 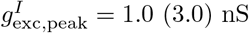

GABA peak conductance on E cells: 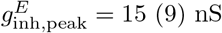

AMPA rise time on I cells: 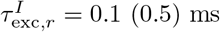

Capacitance of I cells: *C_I_* = 50 (100) pF.

Parameters of the peak dendritic current (cf. Eq 2): *μ* = 0.3 (0.0) nA, *σ* = 0.2 (0.75) nA Peak CA3 rate: *r*_0_ = 9 (8) Hz.

Rectangular rate functions for all CA3 neurons: *r*(*t*) = *r*_0_ (Θ(*t* − (*t*_0_ − *σ*)) − Θ(*t* − (*t*_0_ + *σ*))) with the Heaviside function Θ. The rate function of the CA3 neurons driving all groups except *E*_0_ is centered at *t*_0_ = 56 ms and has width *σ* = 16 ms (black spikes in Fig 9 B top). The rate function of the CA3 neurons driving group *E*_0_ is centered at *t*_0_ = 37 ms and has width *σ* = 4.5 ms (magenta spikes in Fig 9 B top).

#### Higher synaptic I-to-E latency in the replay model

**Fig 29.**
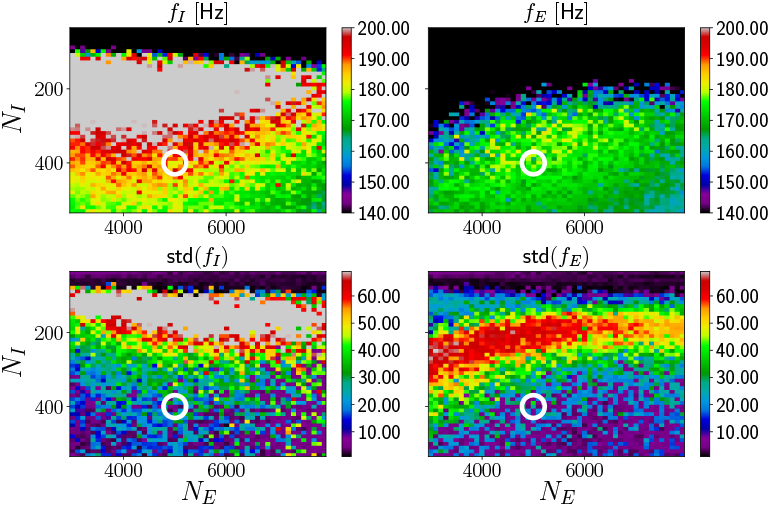
Replay events and ripple oscillations are not robust for longer latency of inhibition. If GABAergic inhibition is received later by the E cells, the replay events and ripples are less robust. The layout and the parameters of the figure are like Fig 10, but the latency of the I-to E-synapses is 1 ms. Replay generates oscillations in the ripple frequency range (upper subpanels). The oscillations in the higher ripple range (about 170 – 190Hz, light green, yellow and red in the upper subpanels) are, however, not robust against changing network realizations: the standard deviation of their frequencies across network realizations is large (lower subpanels). This is because the replay is not robust against changing network realizations: some realizations lead to non-sequential spiking activity with reduced frequency due to increased E cell spiking. For the intermediate and lower ripple frequency range, replay and HFOs are robust.

#### Further alternative concepts of replay

**Fig 30.**
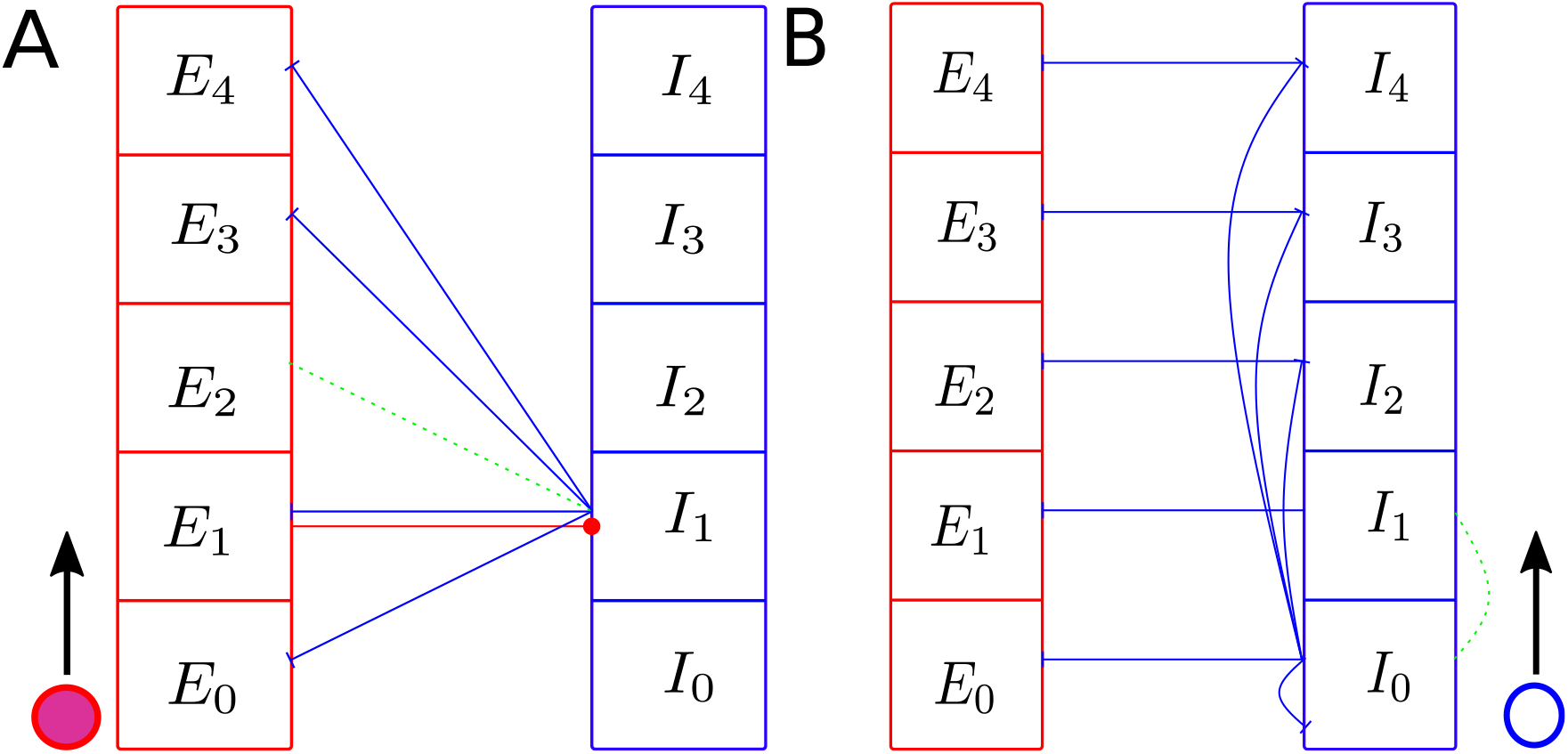
Further connectivity schemes for sequence generation without recurrent excitation. **A:** Scheme for sequence generation by pulse coding of both E and I cells. In contrast to Fig 9 A, the *k*th replay step is in the inhibitory neurons reflected by an activation of group *I_k_*. Each group *E_k_* projects to one group of inhibitory neurons, to *I_k_*, as displayed for *E*_1_. *I_k_* projects to all *E_l_* with *l* ≠ *k* + 1 as displayed for *I*_1_. The green dashed line denotes the absent connections from group *I*_1_ to group *E*_2_. Group *E*_0_ is stimulated to fire first and the replay event progresses along the black arrow: *E*_0_ activates *I*_0_, which inhibits all E cells except those of *E*_1_. Thus, in the next step the group *E*_1_ becomes active and stimulates *I*_1_, which leads to activation of *E*_2_ etc. **B:** Scheme for sequence generation by gap coding of I cells. In contrast to Fig 9 A and panel A of this figure, the sequence generation relies on the inhibitory population alone, no E-to-I connections are necessary. The *k*th replay step is in the inhibitory neurons reflected by a deactivation of group *I_k_*, as in Fig 9 A. There are *K* + 1 inhibitory groups *I*_0_ to *I_K_*. Neurons of group *I*_0_ project to all inhibitory neurons except to those of *I*_1_ (displayed by dashed green line). Similarly, group *I*_1_ does not project to group *I*_2_, but to all other inhibitory groups. Group *I*_1_ thus receives projections from groups *I*_1_ to *I_K_*, group *I*_2_ receives projections from groups *I*_0_ and *I*_2_ to *I_K_* and so on. Every inhibitory group gets projections from *K* inhibitory groups (in the figure: *K* = 4). In the first replay step, group *I*_0_ is silent, whereas all other groups fire. In the next step, every group *I_l_*, *l* > 0, except *I*_1_ receives the same amount of inhibition from only *K* − 1 groups, namely from the previously active groups *I*_1_ to *I*_l−2_ and *I_l_* to *I_K_*, which project to it. The input from *I*_0_ is missing, since it was not active. *I*_1_ receives more inhibition than the other groups because all its *K* inhibitory presynaptic groups were active in the previous step (the silence of *I*_0_ does not reduce *I*_1_’s inhibitory input, since *I*_0_ does not project to *I*_1_). Thus, after the zeroth cycle, group *I*_1_ is silent while the other groups are active. In the next cycle, group *I*_2_ will be silent because it received more inhibition than the other groups and so on. We note that *I*_0_ also gets inhibitory input from *K* groups in each step because it has no predecessor group not inhibiting *I*_0_ but all other groups; its neurons should thus receive a compensatory excitatory input throughout sequence generation to be active after the zeroth step. The E cells in group *E_k_* can be entrained in this scheme by disinhibition, via inhibitory connections from group *I_k_* to *E_k_*.

## Acknowledgments

We thank Avleen Sahni for studying oscillation generation by a network model related to model 1 in her Master’s thesis. We thank Sven Goedeke, Felipe Yaroslav Kalle Kossio, Marcel Stimberg, György Buzsáki and Martin Both for fruitful discussions.

